# Rad51 and Mitotic Function of Mus81 are Essential for Recovery from Low-Dose of Camptothecin in the Absence of the Wrn Exonuclease

**DOI:** 10.1101/399881

**Authors:** Francesca Antonella Aiello, Anita Palma, Eva Malacaria, Li Zheng, Judith L. Campbell, Binghui Shen, Annapaola Franchitto, Pietro Pichierri

## Abstract

Stabilisation of the stalled replication fork is crucial to prevent excessive fork reversal or degradation, which can undermine genome integrity. The WRN protein is a human RecQ helicase that participates in the processing and recovery of perturbed replication forks. WRN is unique among the other human RecQ family members to possess exonuclease activity. However, the biological role of the WRN exonuclease is poorly defined, and little is known about an involvement in the response to perturbed replication. Recently, the WRN exonuclease has been linked to protection of stalled forks from MRE11-dependent degradation in response to clinically-relevant nanomolar doses of the Topoisomerase I inhibitor camptothecin. Alternative processing of perturbed forks has been associated to chemoresistance of BRCA-deficient cancer cells, thus, we used WRN exonuclease-deficiency as a model to investigate the fate of perturbed replication forks undergoing degradation, but in a BRCA wild-type condition. We find that, upon nanomolar doses of camptothecin, loss of WRN exonuclease stimulates fork inactivation and accumulation of parental gaps, which engages RAD51. Such alternative mechanism affects reinforcement of CHK1 phosphorylation and causes persistence of RAD51 during recovery from treatment. Notably, in WRN exonuclease-deficient cells, persistence of RAD51 correlates with elevated mitotic phosphorylation of MUS81 at Serine 87, which is essential to avoid accumulation of mitotic abnormalities. Altogether, these findings indicate that aberrant fork degradation, in the presence of a wild-type RAD51 axis, stimulates RAD51-mediated post-replicative repair and engagement of the MUS81 complex to limit genome instability and cell death.

**AUTHOR SUMMARY:** Correct progression of the molecular machine copying the chromosomes is threatened by multiple causes that induce its delay or arrest. Once the replication machinery is arrested, the cell needs to stabilise it to prevent DNA damage. Many proteins contribute to this task and the Werner’s syndrome protein, WRN, is one of them.

Defining what happens to replication machineries when they are blocked is highly relevant. Indeed, destabilised replication machineries may form upon treatment with anticancer drugs and influence the efficacy of some of them in specific genetic backgrounds. We used cells that lack one of the two enzymatic functions of WRN, the exonuclease activity, to investigate the fate of destabilised replication machineries. Our data show that they are handled by a repair pathway normally involved in fixing DNA breaks but, in this case, recruited to deal with regions of the genome that are left unreplicated after their destabilisation. This alternative mechanism involves a protein, RAD51, which tries to copy DNA from the sister chromosome. In so doing, however, RAD51 produces a lot of DNA interlinking that requires upregulation of a complex, called MUS81/EME1, which resolves this interlinking prior cell division and prevents accumulation of mitotic defects and cell death.

## INTRODUCTION

The response to perturbed replication is crucial for the maintenance of genome integrity [1–5]. In humans, the proper handling of perturbed replication forks is also linked to cancer avoidance and many proteins involved in this process act also as onco-suppressors [3–6]. The importance of dealing correctly with perturbed replication forks is also demonstrated by the existence of several human genetic diseases caused by mutations in factors involved in sensing, processing and recovering replication forks [7].

The Werner’s syndrome protein (WRN) is one of the key factors of the response to perturbed replication and participates directly to the checkpoint in S-phase [8,9]. *WRN* is mutated in the genetic disease Werner’s syndrome (WS), which is characterized by cancer predisposition and premature aging [9], and its loss confers sensitivity to several DNA-damaging agents inducing replication stress [8,10]. From an enzymatic point of view, WRN is both a DNA helicase and exonuclease; however, while its helicase activity has been linked to processing of reversed or collapsed replication forks [2,9], little is known about the biological relevance of the exonuclease activity. We recently reported that the exonuclease activity of WRN is involved in protecting replication forks that are perturbed by treatment with the Topoisomerase I poison Camptothecin (CPT) in the nanomolar range of concentration [11]. Exposure to low doses of CPT, as opposed to high doses, does not induce DSBs in the short-term but stimulates greatly formation of reversed forks [12,13]. Reversed replication forks are versatile yet vulnerable structures and several proteins participate in their stabilisation [14–16]. Two proteins, BRCA2 and RAD51, are the most crucial for the stabilisation of reversed forks [14,16,17]. Thus, cells depleted of each of these two proteins have been used as a prototypical model to assess the consequences of inaccurate handling of reversed forks. However, BRCA2 and RAD51 may also participate in DNA repair, which may be used to fix damage generated by fork instability [17–19]. Loss of WRN exonuclease determines a rapid MRE11-dependent degradation of the nascent strand, most likely after fork reversal, and affects correct replication recovery [11]. However, cells expressing the exonuclease-dead WRN retain ability to restart replication and are not overtly sensitive to low doses of CPT, suggesting that alternative mechanisms can be activated as a back-up. Since nanomolar doses of CPT are clinically-relevant in cancer therapy, cells expressing a catalytically-inactive WRN exonuclease can be used as a model to investigate the fate of CPT-perturbed replication forks undergoing pathological degradation but in a BRCA2-RAD51 wild-type background.

Here, we report that, upon prolonged exposure to nanomolar dose of CPT, loss of WRN exonuclease channels replication forks through a pathological RAD51-dependent mechanism that makes perturbed replication forks resistant to breakage. However, engagement of RAD51 and persisting RAD51 foci make WRN exonuclease-deficient cells reliant on the mitotic function of the MUS81 complex, which mitigates mitotic abnormalities deriving from accumulation of RAD51-dependent intermediates. Furthermore, our data suggest that enhanced accumulation of ssDNA and recruitment of RAD51 interfere with correct activation of CHK1, which provides a positive feedback to the formation of nascent ssDNA.

## RESULTS

### Loss of WRN exonuclease activity leads to a persisting and unusual formation of nascent ssDNA which compromises formation of DSBs in response to a low-dose of camptothecin

Treatment with nanomolar concentrations of CPT does not induce DSBs immediately but initially stimulates fork reversal [12]. Under such conditions, loss of WRN exonuclease activity results in the rapid (< 2h) degradation of both nascent strands by the MRE11-EXO1 nucleases [11]. In this regard, WRN exonuclease deficiency is a useful model to determine what happens to nuclease-targeted forks after prolonged treatment with low dose of CPT. This prompted us to analyse the processing of perturbed replication forks beyond 2h of treatment.

We first examined the presence of nascent ssDNA as a sign of fork degradation by the native IdU assay. As expected, WS cells expressing the exo-dead WRN protein (WS^E84A^) showed less nascent ssDNA then the corrected wild-type counterpart (WS^WT^) at 1h of treatment (Fig. 1A). Surprisingly, the amount of nascent ssDNA in WRN exonuclease-deficient cells increased greatly over time and largely exceeded the wild-type level at 4h, while it increased only slightly in cells expressing the wild-type WRN (Fig. 1A). Since nascent strand degradation is MRE11-dependent in the absence of the WRN exonuclease while it is DNA2-dependent in wild-type cells (refs), we next examined if chemical inhibition of those nucleases might reduce the idiosyncratic accumulation of ssDNA detected at 4h of treatment with nanomolar CPT in WS^E84A^ cells. Mirin treatment, which inhibits MRE11, barely reduced ssDNA detected at 4h of treatment in wild-type cells (Fig. 1B). Surprisingly, Mirin did not decrease the formation of ssDNA in WRN exonuclease-deficient cells but rather increased it even further (Fig. 1B). Inhibition of DNA2 by the small-molecule inhibitor C5 [20] increased formation of nascent ssDNA in wild-type but not in WRN exonuclease-deficient cells (Fig. 1B). In contrast, concomitant inhibition of DNA2 and MRE11 was ineffective in modulating nascent ssDNA formation in wild-type cells while decreased its level in cells expressing the exo-dead form of WRN (Fig. 1B). Of note, ssDNA derived from end-resection of DSBs induced by a micromolar dose of CPT was efficiently reduced by DNA2 inhibitor C5 (Fig. S1A, B), providing a functional proof of the inactivation of the nuclease activity by the C5 compound in our cell model. These results indicate that different sets of nucleases are involved in the degradation of nascent ssDNA in WRN exonuclease-deficient cells when treatment is prolonged.

**Figure 1.**
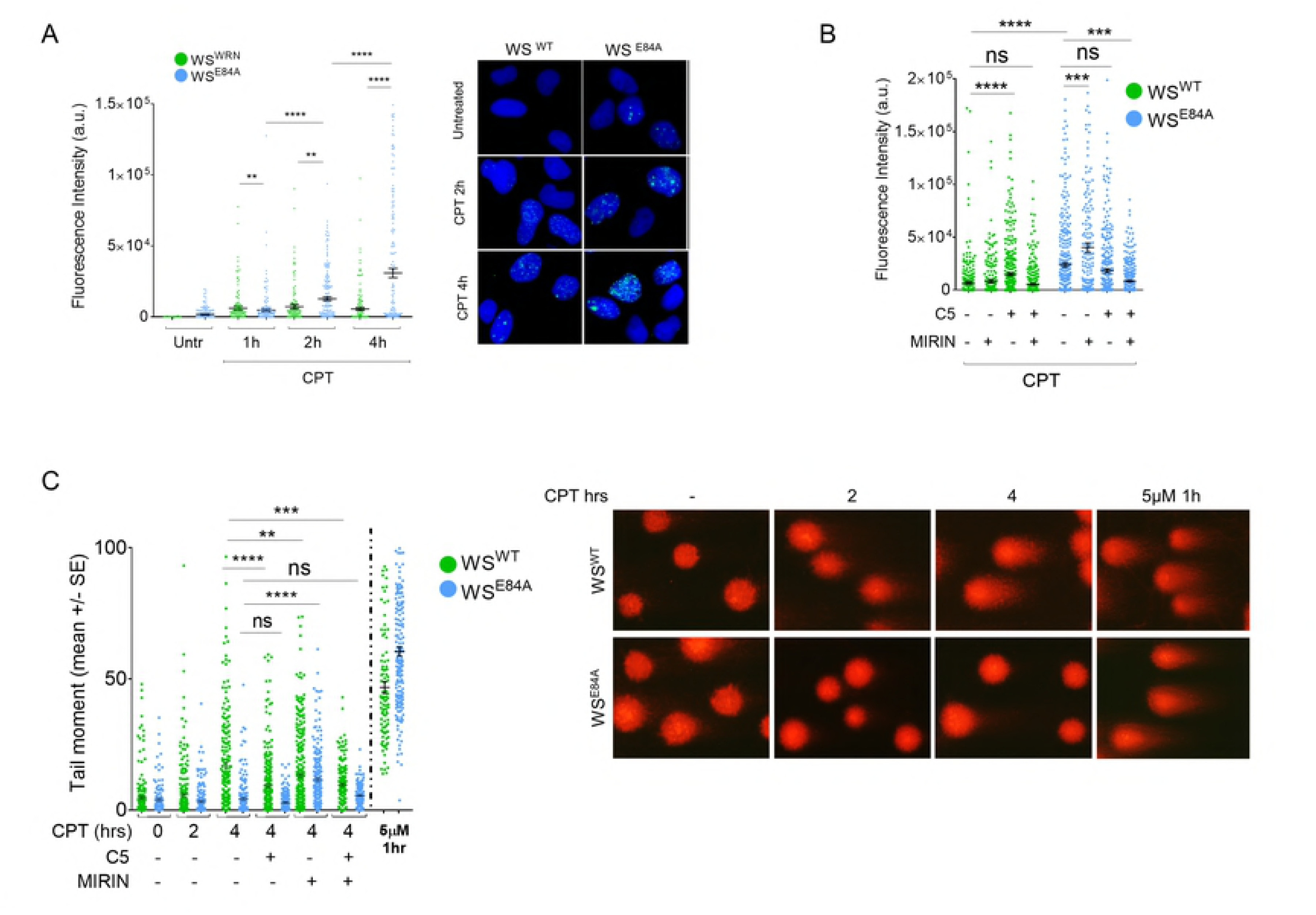
Loss of WRN exonuclease activity leads to formation of nascent ssDNA which compromises formation of DSBs in response to a low-dose of camptothecin. (A) Evaluation of ssDNA by anti-IdU immunofluorescence under non-denaturing condition. Nascent DNA was pre-labelled for 15 min with IdU before treatment and labelling remained during treatment with CPT. Dot plots show the mean intensity of ssDNA staining for single nuclei from cells expressing the wild-type (WS^WT^) or the exo-dead form of WRN (WS^E84A^). Cells were either left untreated or challenged with 50 nM CPT for increasing periods, as indicated. The intensity of the anti-IdU immunofluorescence was measured in at least 200 nuclei from three independent experiments. Values are represented as means ±SE. Representative images of ssDNA labelling are shown. (B) Evaluation of nascent ssDNA in cells treated with nuclease inhibitors. Cells were treated with Mirin, C5 or both 30 min before IdU labelling and 45 min before CPT treatment for 4 h, and then subjected to the ssDNA assay. The graph shows the mean intensity of IdU fluorescence measured from two independent experiments (n=200), data are presented as mean ±SE. Statistical analysis in A and B was performed by the Mann–Whitney test (\*\**P* < 0.1; \*\*\**P <* 0.01; \*\*\*\**P <* 0.001) (C) Analysis of DSB accumulation by the neutral Comet assay. Cells were treated or not with CPT 50 nM for the indicated time, or with 5μM CPT (high-dose) for 1 h, and then subjected to the neutral Comet assay. Where indicated, cells were pre-treated with Mirin, C5 or both. In the graph, data are presented as mean tail moment ± SE from two independent experiments (ns = not significant; \*\**P <* 0.1; \*\*\**P <* 0.01; \*\*\*\**P <* 0.001; ANOVA test). Representative images from the neutral Comet assay are shown.

DNA breakage can occur even in response to low-doses of CPT if treatment is sufficiently prolonged [12,13]. Thus, to further investigate the origin of the late nascent ssDNA in the WRN exonuclease-deficient cells and the role of the different nucleases, we analysed the presence of DSBs after treatment with nanomolar CPT by neutral Comet assay. As shown in Figure 1C, treatment with 50nM CPT for 4h is able to induce some DSBs in wild-type cells, although they are very low compared with those generated by the 5μM reference dose. In contrast, no DSBs were detected in WRN exonuclease-deficient cells after treatment with the low-dose of CPT, even if they were readily seen in response to the high-dose of the drug (Fig. 1C). Interestingly, pre-treatment with Mirin, which enhances ssDNA in WS^E84A^ cells (Fig. 1B), resulted in DSBs (Fig. 1C). In contrast, formation of DSBs was unaffected in wild-type cells by Mirin while it was reduced by the DNA2 inhibitor (Fig. 1C). Interestingly, inhibition of both MRE11 and DNA2, which decreases ssDNA formation in WS^E84A^ cells (Fig. 1B), reduced significantly the DSBs generated by Mirin (Fig. 1C). Concomitant inhibition of MRE11 and DNA2 also reduced DSBs in wild-type cells (Fig. 1C). The analyses of DSBs and ssDNA suggest that the increase in nascent ssDNA induced by MRE11 inhibition in WRN exonuclease-deficient cells derives from end-resection of DSBs. In contrast, the reduction of nascent ssDNA induced by concomitant treatment with C5 and Mirin would indicate that DNA2 and MRE11 are involved in the formation of the late nascent ssDNA observed in WS^E84A^ cells independently of detectable DSBs. Finally, we tested whether MUS81 was required for late DSB formation [21,22]. As shown in Fig. S1C, DSBs derived from prolonged treatment with a low-dose of CPT in wild-type cells are not affected by depletion of MUS81.

Collectively, our results indicate that loss of the WRN exonuclease leads to accumulation of nascent ssDNA when treatment with nanomolar CPT is prolonged beyond 2h and that it makes cells refractory to formation of DSBs by CPT. Our data also suggest that the late accumulation of nascent ssDNA in WS^E84A^ cells is related to the activity of multiple nucleases.

### Loss of WRN exonuclease stimulates engagement of RAD51 after CPT-induced fork perturbation

In WRN exonuclease-deficient cells, late reappearance and accumulation of nascent ssDNA together with absence of DSBs after CPT might correlate with engagement of an alternative or aberrant fork processing mode over time. Extended regions of ssDNA are a substrate for RAD51 during recombination, thus, we investigated whether loss of WRN exonuclease could affect recruitment of RAD51 after treatment with CPT.

To evaluate recruitment of RAD51, we first prepared chromatin from cells treated with nanomolar CPT and determined the amount of RAD51 present by Western blotting. In wild-type cells, RAD51 barely increased its chromatin association after CPT treatment (Fig. 2A). Expression of the exo-dead WRN, however, greatly increased the amount of RAD51 in chromatin both in untreated cells and CPT-treated cells. Treatment with CPT led to a minimal increase over untreated (about 20%; Fig. 2A). As a further control, we also measured the level of RPA32, a subunit of the RPA heterotrimer that binds to ssDNA, in chromatin. RPA32 increased after CPT in wild-type cells (Fig. 2A). In contrast, the amount of RPA32 in chromatin was low in the absence of the WRN exonuclease and did not show any increase after treatment (Fig. 2A). RAD51 participates in multiple pathways [23,24] and chromatin recruitment may reflect such pleiotropy of roles. To further verify the increased recruitment of RAD51 in cells expressing the exo-dead WRN, we performed a quantitative immunofluorescence analysis of the RAD51 foci (Fig. 2B). We evaluated the number of foci by analysing the intensity of fluorescence in the RAD51 foci-positive nuclei. Consistent with biochemical data, loss of WRN exonuclease increased recruitment of RAD51 (Fig. 2B). Moreover, CPT treatment led to a higher number of RAD51 foci in WRN exonuclease-deficient cells compared with the wild-type (Fig. 2B)

**Figure 2.**
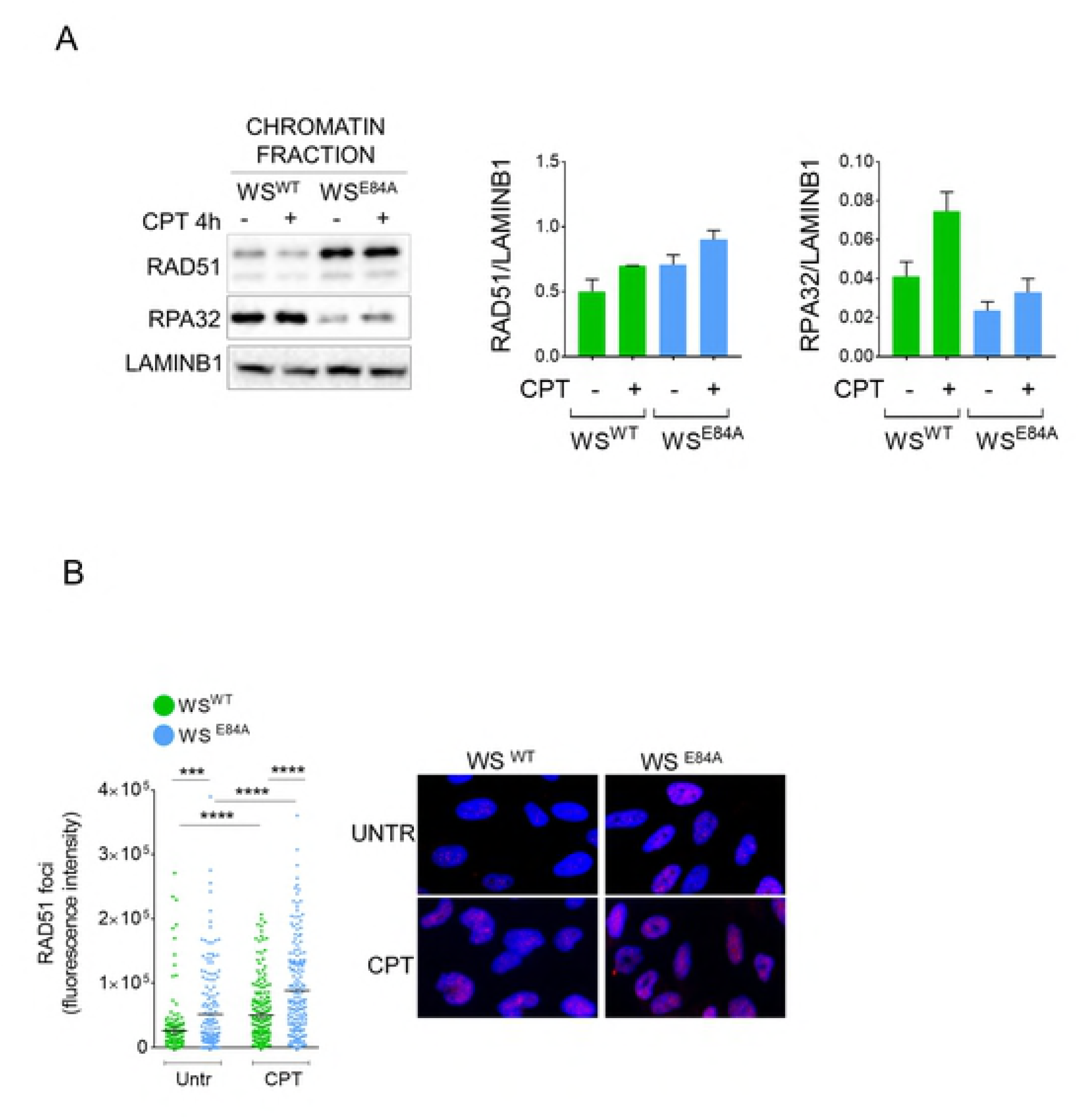
Loss of WRN exonuclease stimulates engagement of RAD51 after CPT. (A) WB analysis of chromatin association of RAD51 and RPA32 in wild-type (WS^WT^) and in cells expressing the exo-dead mutant form of WRN (WS^E84A^). Cells were treated or not with CPT for 4 h. LaminB1 was used as loading control. The blot is representative of three replicates. The graphs show the quantification of the amount of RAD51 or RPA32 normalised against LaminB1 (mean±SE). (B) Quantitative immunofluorescence analysis of RAD51 foci in WS^WT^ and WS^E84A^ cells. Cells were treated with CPT 50nM for 4h, triton-extracted and subjected to RAD51 immunostaining. Graph shows the intensity of RAD51 immunostaining for each cell with scorable foci (n>3). Values are presented as means ± SE (*** *P* < 0.01;**** *P* < 0.001; Mann–Whitney test). Representative images are shown.

These results suggest that, in the absence of the WRN exonuclease, RAD51 takes over the normal mechanism handling CPT-perturbed replication forks, perhaps to provide a backup mechanism for recovery.

To test if RAD51 participated in replication fork restart, we analysed whether, in cells expressing the exo-dead WRN, RAD51 chromatin levels remained elevated also during recovery from CPT. To this end, we treated cells with CPT and allowed them to recover for 2 and 4h prior to preparing chromatin fractions. As shown in Figure 3A, while RAD51 levels, as determined by Western blotting, tended to remain low during recovery in wild-type cells, they remained elevated in WRN exonuclease-deficient cells. Of note, the amount of chromatin-bound RPA32 and RPA70, two subunits of trimeric RPA, remained low in WS^E84A^ cells while it increased greatly during recovery in cells expressing the wild-type WRN (Fig. 3A). To confirm this result, we performed quantitative immunofluorescence analysis of the formation of RAD51 foci during recovery from CPT. A reduction of RAD51 foci was observed during recovery from CPT in wild-type cells (Fig. 3B). In contrast, the number of RAD51 foci increased in WRN exonuclease-deficient cells (Fig. 3B), confirming that a RAD51-dependent pathway is over-activated in response to prolonged treatment with low-dose of CPT in these cells.

**Figure 3.**
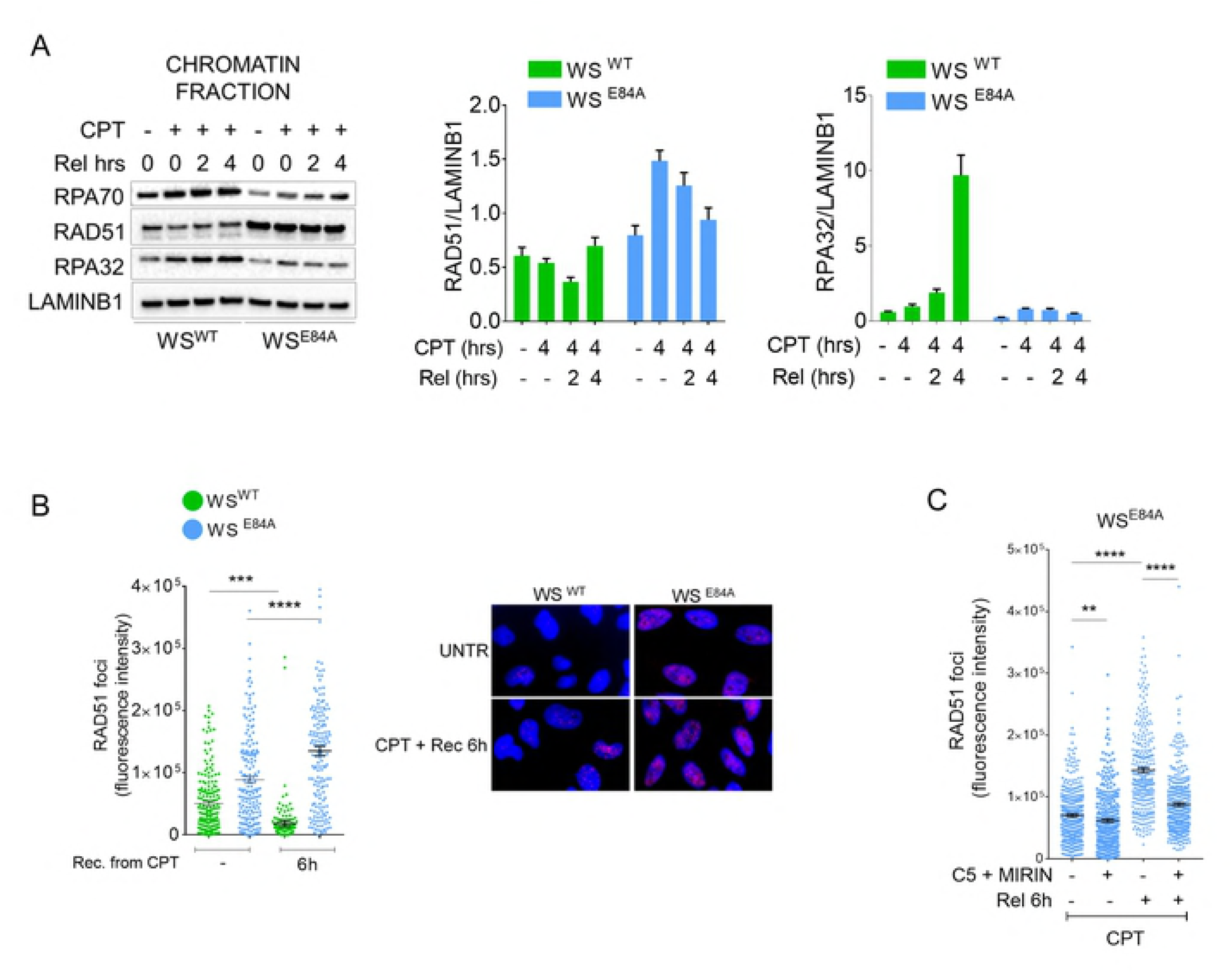
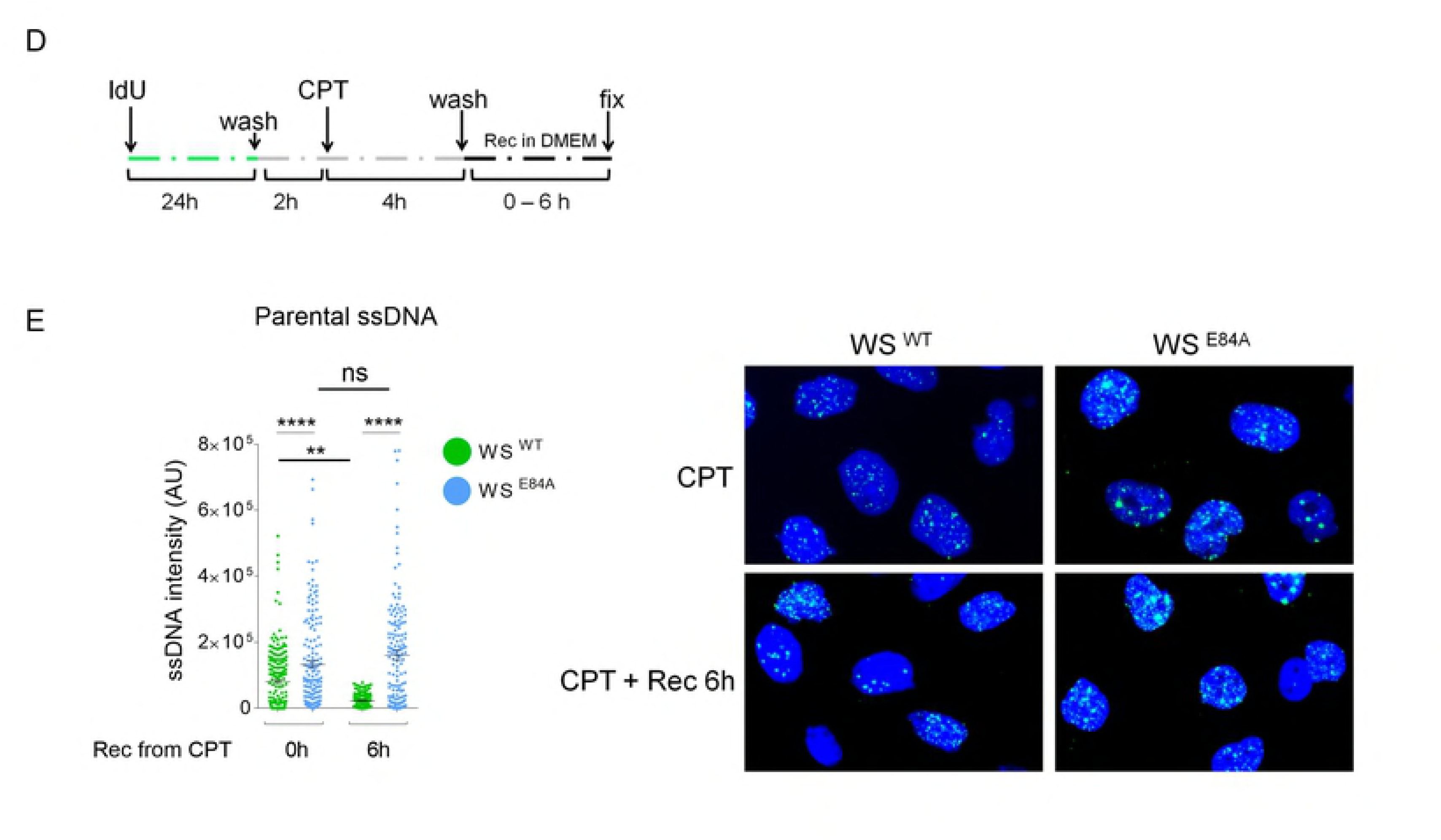
RAD51 recruitment persisted during recovery from CPT in the absence of the WRN exonuclease. (A) WB analysis of chromatin association of RAD51, RPA70 and RPA32 in wild-type (WS^WT^) and in cells expressing the exo-dead mutant form of WRN (WS^E84A^). Cells were treated or not with 50nM CPT for 4h followed by recovery as indicated. LaminB1 was used as loading control. The blot is representative of three replicates. The graphs show the quantification of the amount of RAD51 or RPA32 normalised against LaminB1 (mean±SE). (B) Quantitative immunofluorescence analysis of RAD51 foci in WS^WT^ and WS^E84A^ cells. Cells were treated with CPT 50nM for 4h and recovered or not as indicated. Graph shows the intensity of RAD51 immunostaining for each cell with scorable foci (n>3). Values are presented as means ± SE (*** *P* < 0.01; **** *P* < 0.001; Mann– Whitney test). Representative images are shown. (C) Quantitative immunofluorescence analysis of RAD51 foci in WS^WT^ and WS^E84A^ cells pre-treated with the nuclease inhibitors. Cells were pre-treated with the indicated inhibitors prior to be challenged with CPT 50nM for 4h and recovered or not as indicated. Graph shows the intensity of RAD51 immunostaining for each cell with scorable foci (n>3). Values are presented as means ± SE (** *P* < 0.1; **** *P* < 0.001; Mann–Whitney test). (D-E) analysis of parental ssDNA. Parental DNA was labelled with IdU as indicated in the experimental scheme (D). (E) The graph shows the amount of parental ssDNA calculated as mean intensity of IdU fluorescence measured from two independent experiments (n=200), data are presented as mean ±SE. Statistical analysis was performed by the Mann–Whitney test (\*\**P* < 0.1; \*\*\*\**P <* 0.001). Representative images are shown.

The persistently-high levels of RAD51 observed in the absence of the WRN exonuclease even during recovery prompted use to determine if they were correlated with the increased formation of nascent ssDNA. Having demonstrated that concomitant inhibition of MRE11 and DNA2 restores wild-type levels of nascent ssDNA in WRN exonuclease-deficient cells (Fig. 1C), we analysed RAD51 focus-forming activity after pre-treatment of cells with C5 and Mirin. As expected, RAD51 recruitment in foci was elevated in WRN exonuclease-deficient cells after treatment and even more during recovery (Fig. 3C). Interestingly, concomitant pre-treatment with C5 and Mirin significantly reduced the formation of RAD51 foci in WS^E84A^ cells (Fig. 3C), suggesting that it depends on late accumulation of nascent ssDNA observed in the absence of WRN exonuclease.

The elevated levels of RAD51 during recovery might be related to the presence of under-replicated DNA that requires recombination to be replicated or repaired [25]. Thus, we analysed whether cells recovering from CPT treatment presented under-replicated regions of DNA. Since under-replication is expected to leave regions of ssDNA in the parental strand (parental gaps) behind perturbed forks, we performed native IdU assay after a 24h treatment with IdU to label all parental DNA prior to add CPT and perform recovery (Fig. 3D). As shown in Figure 3E, little parental ssDNA was detectable in wild-type cells after treatment and its amount decreased substantially during recovery. In WS^E84A^ cells, however, parental ssDNA was higher at the end of treatment and remained elevated during recovery (Fig. 3E). Interestingly, the increased amount of parental ssDNA paralleled that of RAD51 foci, suggesting that RAD51 may be recruited to deal with under-replicated regions.

The presence of under-replicated DNA and the increased recruitment of RAD51 might indicate that perturbed replication forks get inactivated upon prolonged exposure to CPT in the absence of the WRN exonuclease. Thus, we carried out a single-molecule analysis of DNA replication by the DNA fiber assay (Fig. 4A). As observed previously [12,13], treatment with nanomolar CPT did not induce fork arrest but rather a delay in fork progression (Fig. S2). Indeed, the length of the CldU tract is increased during extended periods of treatment. Of note, length of the CldU tract was similar between WS^WT^ and WS^E84A^ cells at 1h and 4h of treatment (Fig. S2). When we analysed the ability to replicate after treatment, we found that most of perturbed replication forks remained active after 1h of CPT in wild-type cells, as indicated by the low level of stalled forks (red-only tracks) (Fig. 4B-D). After prolonged treatment, the number of stalled forks increased by around 2-fold, but perturbed forks remained mostly active (Fig. 4B). Notably, the number of stalled forks was higher in WRN exonuclease-deficient cells and around the 50% of the forks got inactivated at 4h of treatment (Fig. 4B-D). Interestingly, in the absence of the WRN exonuclease, treatment with CPT stimulated firing of new origins (green-only tracks), which increased with time of treatment (Fig. 4C, D). In contrast, limited new origin firing was detectable in wild-type cells (Fig. 4C, D). Surprisingly, and in spite of the elevated levels of RAD51 in WS^E84A^ cells, neither the percentage of inactive forks nor that of new origins was affected by inhibition of RAD51 during recovery (Fig. 4B-D). In contrast, we observed that both fork inactivation and firing of new origins were increased by inhibition of RAD51 in wild-type cells (Fig. 4B-D).

**Figure 4.**
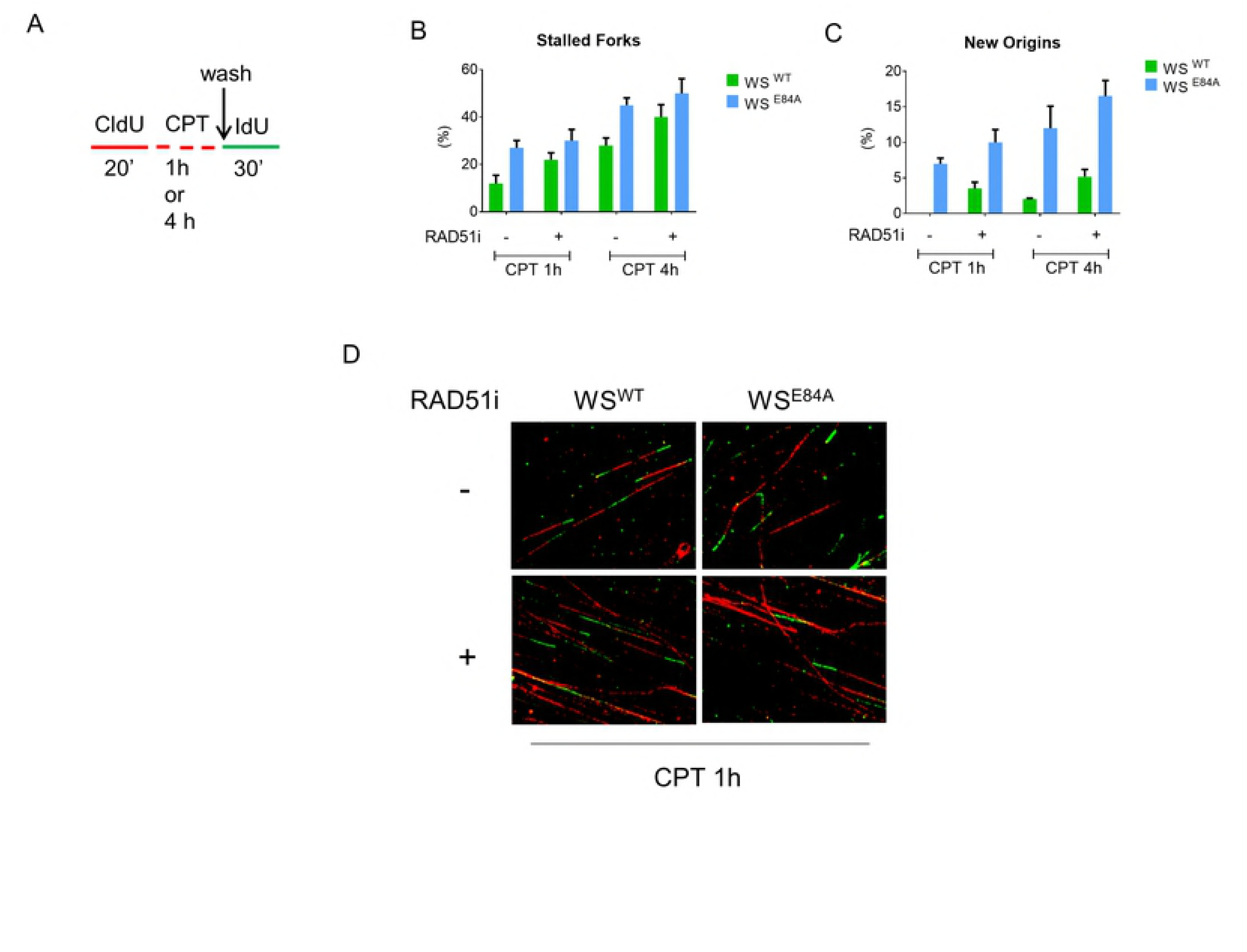
RAD51 inhibition does not impair replication fork recovery following treatment with low dose of CPT. (A) Experimental scheme of dual-labelling replication assay with DNA fibres. Red tract: CldU; Green tract: IdU. (B) The graph shows the average number of stalled forks (red only tracts opposed to the active forks marked as red+green tracks) after recovery from 50nM CPT treatment. Where indicated, RAD51 inhibitor (B02; RAD51i) was added to cultures together with CPT and during the IdU pulse. Data are presented as mean ± SE. One-hundred IdU-positive tracts were analysed in each experimental point (n=2). (C) The graph shows the average number of new origins (green only tracts) after recovery from 50nM CPT treatment. Where indicated, RAD51 inhibitor (B02; RAD51i) was added to the cultures together with CPT and during the IdU pulse. Values are presented as mean ± SE. In B and C statistical analysis was performed by Anova test. (D) Representative DNA fibres fields are shown in the images.

These results suggest that RAD51 is not required to promote replication recovery but rather to promote repair of parental gaps left behind inactive forks after resumption of synthesis when the exonuclease activity of WRN is absent.

### Loss of WRN exonuclease reduces activation of CHK1

Our data indicate that loss of the WRN exonuclease results in accumulation of ssDNA and RAD51 accompanied by a concomitant decrease of RPA. Since RPA-coated ssDNA is required for checkpoint signalling upon replication fork perturbation, we investigated whether the functionality of the WRN exonuclease might also affect activation of the replication checkpoint in response to nanomolar concentration of CPT. As a readout of the activation of the ATR-dependent checkpoint response, we analysed phosphorylation of CHK1 at S345 by Western blotting. In wild-type cells, treatment with a nanomolar dose of CPT induced a time-dependent phosphorylation of CHK1 which is also readily observed in cells expressing the helicase-dead form of WRN (WS^K577M^) (Fig. 5A). In contrast, CPT-induced phosphorylation of CHK1 was reduced in cells expressing the exonuclease-dead mutant of WRN, and this phenotype was more evident at 4 and 6h of treatment (Fig. 5A). The requirement of the WRN exonuclease for correct CHK1 phosphorylation was specific for the low-dose CPT treatment as it was not observed after 5μM of CPT (Fig. S3A).

**Figure 5.**
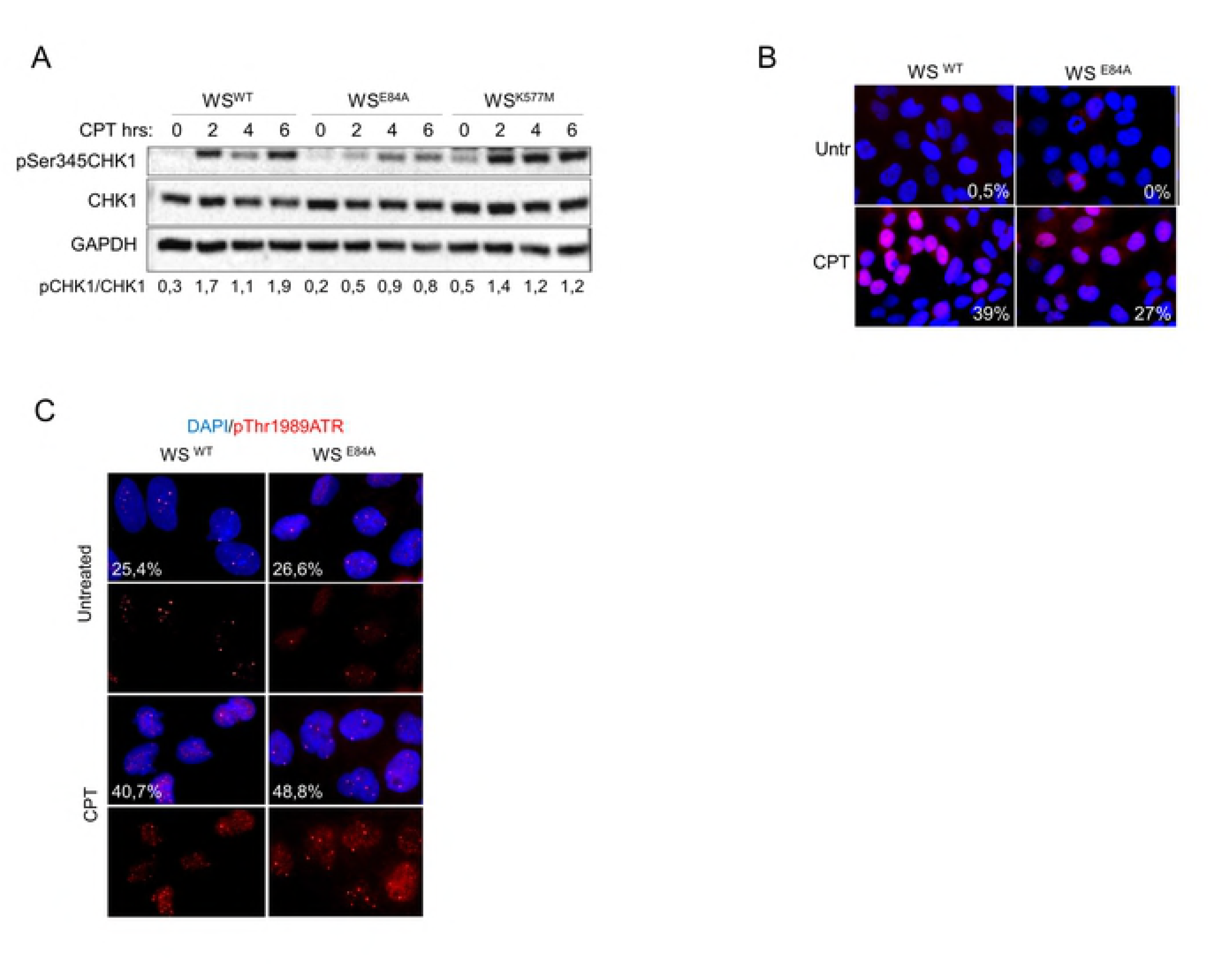
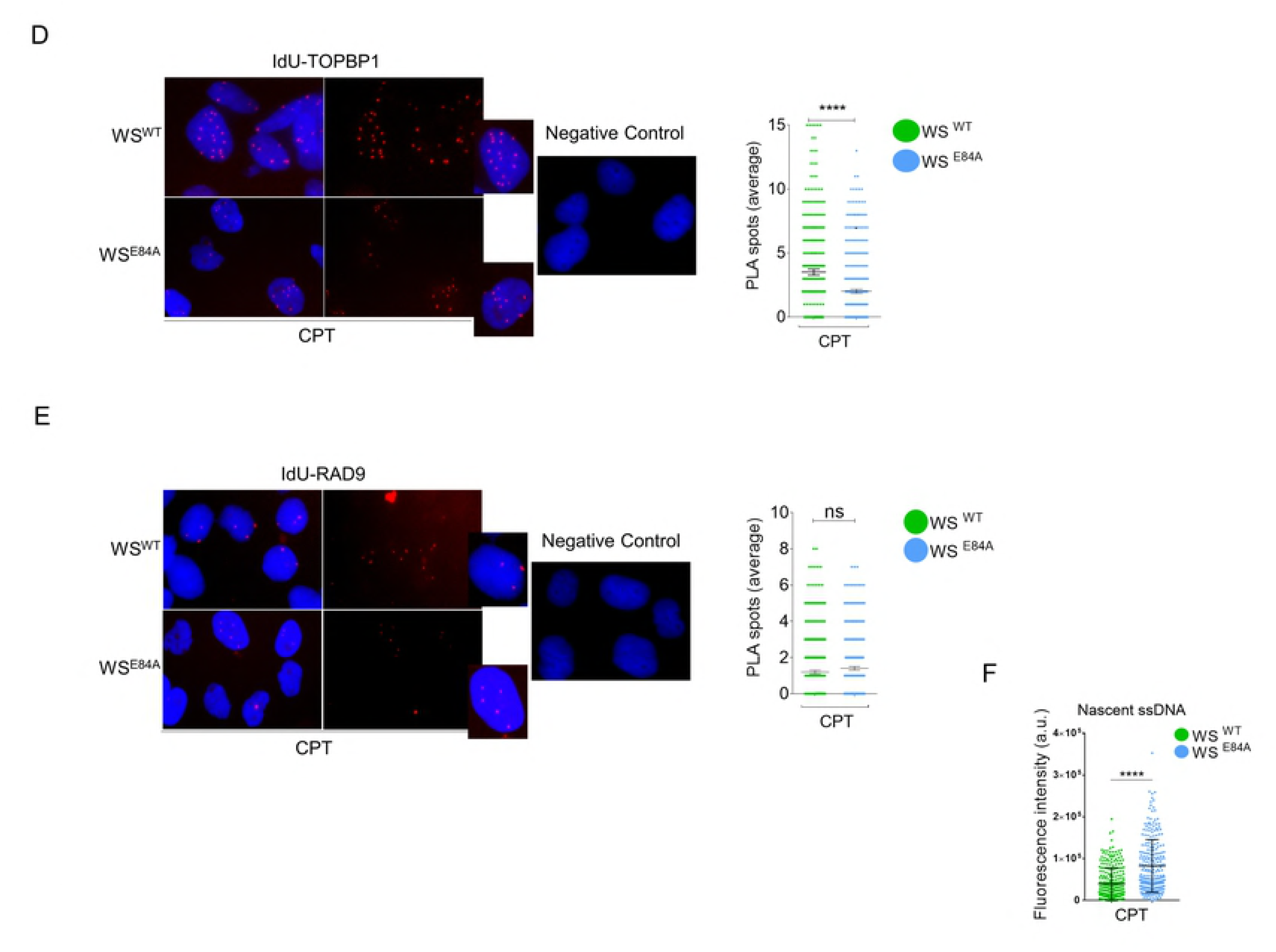
Loss of WRN exonuclease activity affects phosphorylation of CHK1. (A) WB analysis of CHK1 phosphorylation at S345 in wild-type (WS^WT^) and in cells expressing the exo-dead mutant form of WRN (WS^E84A^) or the helicase-dead form (WS^K577M^). Cells were treated or not with 50nM CPT as indicated. Total CHK1 and GAPDH were used as loading controls. The blot is representative of three replicates. Below is reported quantification of p345CHK1 phosphorylation normalised against total CHK1. (B) Immunofluorescence analysis of pS345CHK1 in WS^WT^ and WS^E84A^ cells treated with CPT 50nM for 4h. Numbers in insets represent the mean percentage of pS345CHK1-positive nuclei (n=2; errors are not shown but are < 15% of the mean). (C) Immunofluorescence analysis of pT1989ATR in WS^WT^ and WS^E84A^ treated with CPT 50nM for 4h. Numbers in insets represent the mean percentage of pS345CHK1-positive nuclei (n=2; errors are not shown but are < 15% of the mean). (D-E) Analysis of TopBP1 or RAD9 recruitment at nascent ssDNA by PLA. Nascent strand was labelled with IdU for 15min before cells were treated with 50nM CPT for 4h. PLA was performed under native conditions using anti-IdU to detect nascent ssDNA and anti-TopBP1 or RAD9 to detect the protein. Negative controls are from samples processed with anti-IdU only. The graphs show the number of PLA spots in each nucleus (n=300 from 3 independent replicates). Statistical analysis was performed by the Mann–Whitney test (ns = not significant; \*\*\*\**P <* 0.001). Representative images are shown. (F) Duplicated samples from D-E were analysed for the presence of nascent ssDNA by native anti-IdU detection only. The graph shows the mean intensity of IdU fluorescence measured from two independent experiments (n=200), data are presented as mean ±SE. Statistical analysis was performed by the Mann–Whitney test (\*\*\**P <* 0.01).

To confirm that loss of WRN exonuclease affected CHK1 phosphorylation by an independent assay, we monitored the status of S345 of CHK1 by immunofluorescence. As shown in Fig. 5B, a reduced phosphorylation of CHK1 at S345 was readily detected also by immunofluorescence in WRN exonuclease-deficient cells.

Next, we wanted to analyse whether decreased activation of CHK1 correlated with reduced activation of ATR-dependent signalling. To assess activation of ATR, we monitored phosphorylation of the activating site T1989 by immunofluorescence. Despite the defective phosphorylation of CHK1, ATR was phosphorylated similarly in wild-type cells and in cells expressing the exonuclease-dead form of WRN (Fig. 5C). Since loss of WRN exonuclease affects recruitment of RPA but did not affect activation of ATR, we analysed whether it could influence recruitment of other factors modulating ATR-checkpoint function. As loss of WRN exonuclease leads to accumulation of nascent ssDNA at 4h of CPT (Fig. 1A), we analysed the presence of TopBP1 and RAD9, which associates with TopBP1 [26,27], specifically at nascent ssDNA by our recently described IdU-PLA assay [11]. In parallel, we evaluated the presence of the total amount of ssDNA by IdU assay. Association of TopBP1 or RAD9 with nascent ssDNA was not detected under untreated conditions (data not shown), however, treatment with 50nM CPT for 4h resulted in recruitment of both factors at nascent ssDNA in wild-type cells (Fig. 5D, E). Of note, loss of WRN exonuclease reduced the recruitment of TopBP1 but not that of RAD9 at the nascent ssDNA (Fig. 5D, E) although in these conditions the IdU assay detected 2-times more ssDNA (Fig. 5F).

Loss of the WRN exonuclease leads to accumulation of nascent ssDNA, which is targeted by RAD51 and not by checkpoint factors, possibly resulting in reduced CHK1 phosphorylation. Thus, we investigated whether pre-treatment with the RAD51 inhibitor B02 or treatment before sampling could re-establish a normal CHK1 activation in WRN exonuclease-deficient cells (Fig. S3B). Of note, inhibition of RAD51 further decreased the phosphorylation of CHK1 regardless the way it was added (Fig. S3B). This was surprising but prompted us to evaluate whether reduced CHK1 activation could be implicated in the enhanced accumulation of nascent ssDNA. To test this potential feedback effect, we expressed the S317/S345D CHK1 phosphomimetic mutant [28] in WS^E84A^ cells (Fig. 6A) before evaluating the formation of nascent ssDNA by the IdU assay. As shown in Figure 6B, the phosphomimetic CHK1 was efficiently expressed in the cells and its expression was able to substantially reduce the amount of nascent ssDNA in WRN exonuclease-deficient cells restoring the wild-type levels.

**Figure 6.**
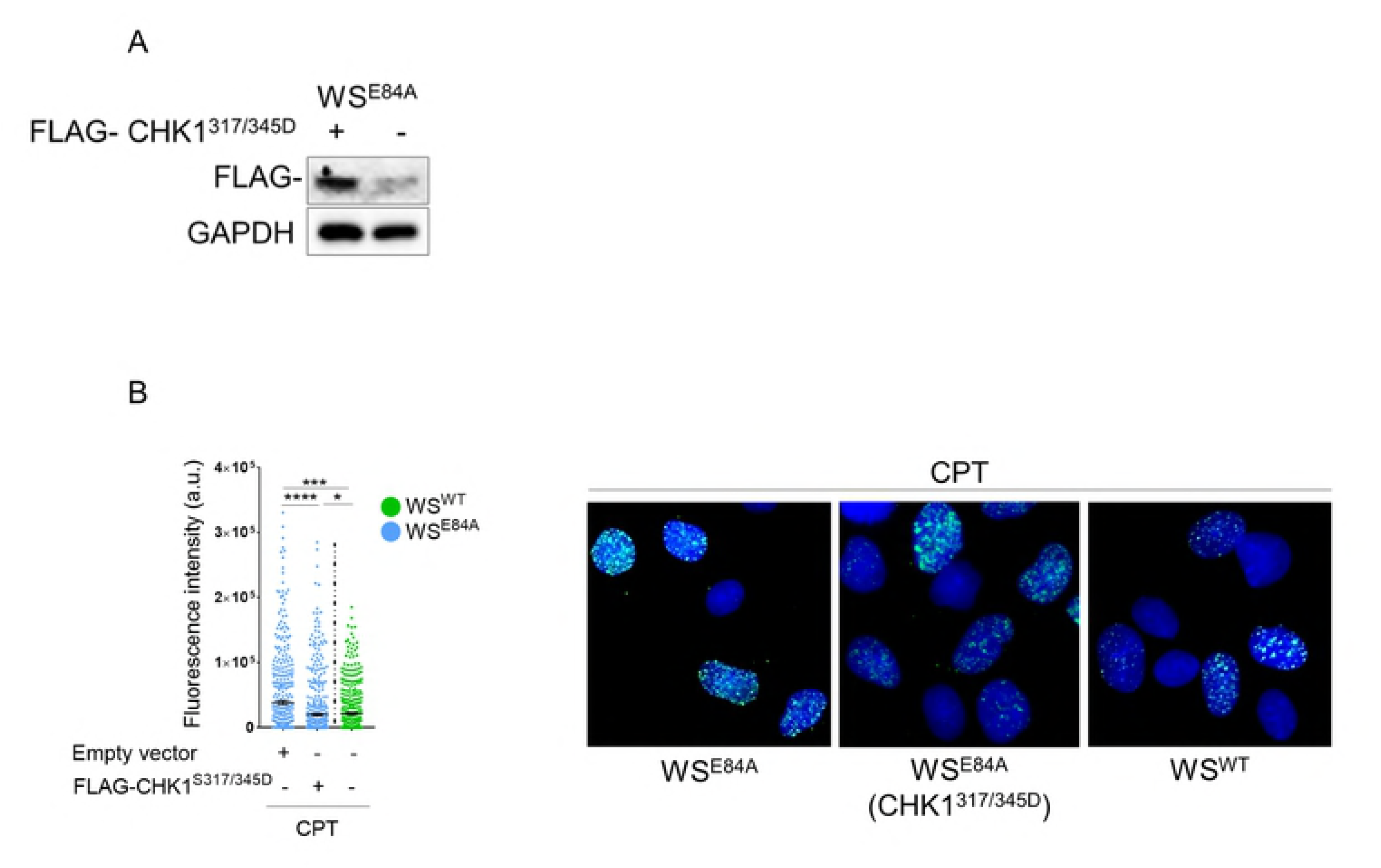
Expression of a phosphomimic CHK1 mutant restores wild-type levels of nascent ssDNA in WRN exonuclease-deficient cells. (A) WB analysis of FLAG-CHK1^317/345D^ expression in WS^E84A^ cells. (B) Evaluation of nascent ssDNA formation. Cells treated with 50nM CPT for 4h were analysed for the presence of nascent ssDNA by native anti-IdU detection. The graph shows the mean intensity of IdU fluorescence measured from two independent experiments (n=200), data are presented as mean ±SE. Statistical analysis was performed by the Mann–Whitney test (\**P <* 0.5; \*\*\**P <* 0.01; \*\*\*\**P <* 0.001). Representative images are shown.

Altogether, our results show that loss of WRN exonuclease activity affects proper activation of CHK1 in response to a low-dose of CPT and that reduced phosphorylation of CHK1 probably correlates with reduced recruitment of checkpoint factors at ssDNA. They also suggest that reduced CHK1 phosphorylation contributes to the accumulation of ssDNA in the nascent strand, possibly as part of a positive feedback loop.

### WRN exonuclease-deficient cells need the mitotic function of MUS81 to counteract mitotic aberration and mis-segregation

Inaccurate processing of perturbed replication forks, elevated under-replicated DNA and RAD51 levels observed in the absence of WRN exonuclease could threaten mitosis because of DNA interlinking as shown in BRCA2-deficient cells [29]. Mitotic resolution of DNA interlinked intermediates involves the MUS81/EME1 complex [30,31]. Thus, we evaluated whether WRN exonuclease-deficient cells accumulated active MUS81 in mitosis by performing immunofluorescence with an antibody directed against the pS87-MUS81, which we have demonstrated to be a readout of the active MUS81/SLX4 complex [32]. In wild-type cells, little MUS81 phosphorylation at S87 was detectable either under unperturbed cell growth or in response to low-dose of CPT (Fig. 7A). In contrast, many pS87-MUS81-positive nuclei were detectable in cells expressing the exonuclease-dead WRN already in unperturbed conditions (Fig. 7A). Notably, in WRN exonuclease-deficient cells, pS87-MUS81 levels were further enhanced by CPT and remained elevated also after recovery (Fig. 7A). Interestingly, and consistent with our previous results [32], phosphorylation of MUS81 never occurred in S-phase cells labelled with EdU (Fig. 7A). To determine whether elevated MUS81 activation correlated with under-replication and enhanced RAD51 recruitment, we asked if inhibition of RAD51 by B02 (ref) reverted the pS87-MUS81 levels in WRN exonuclease-deficient cells. As shown in Figure 7B, inhibition of RAD51 greatly decreased activation of the MUS81 complex after recovery from CPT, as indicated by reduction in pS87-MUS81-positive nuclei.

**Figure 7.**
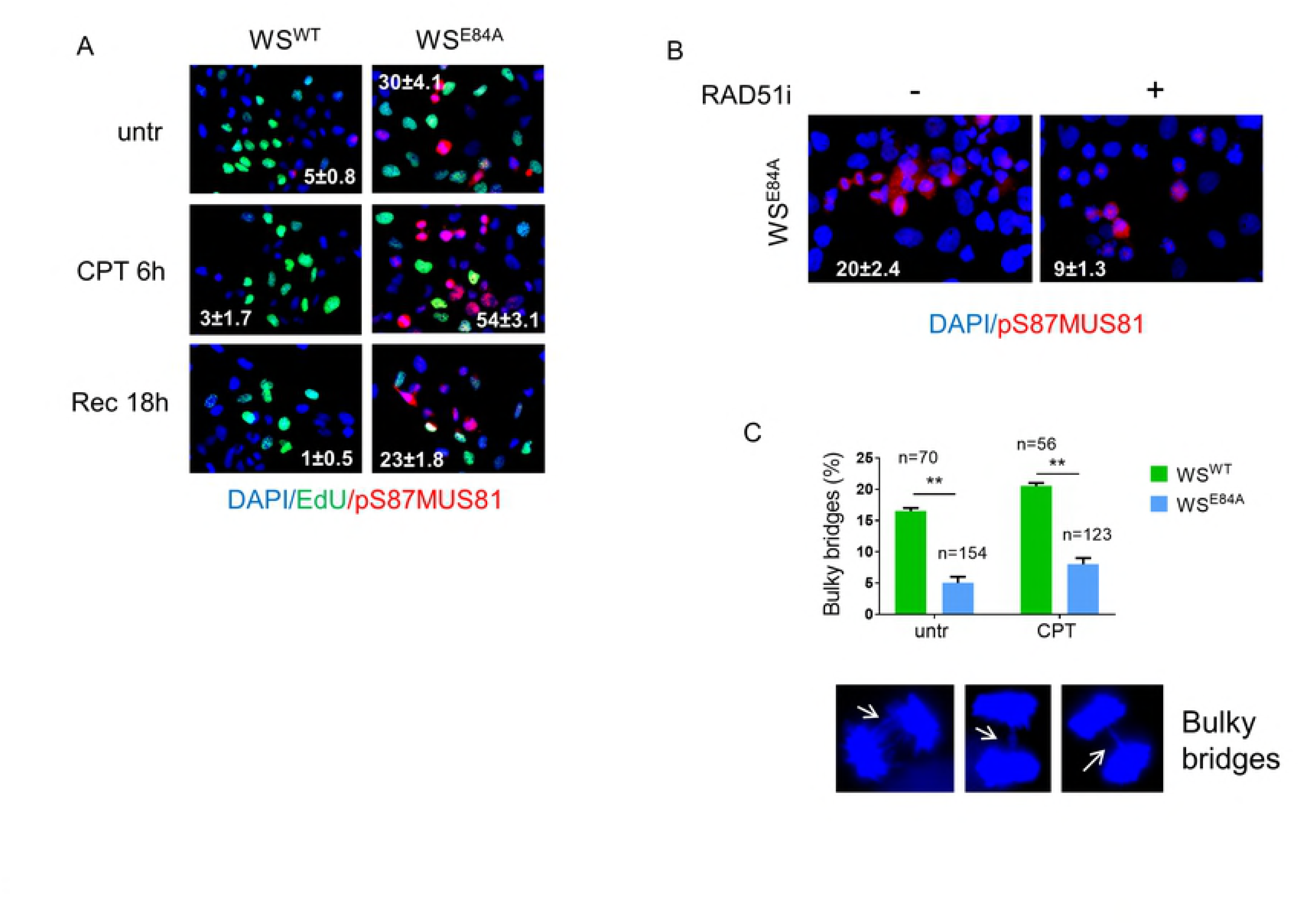
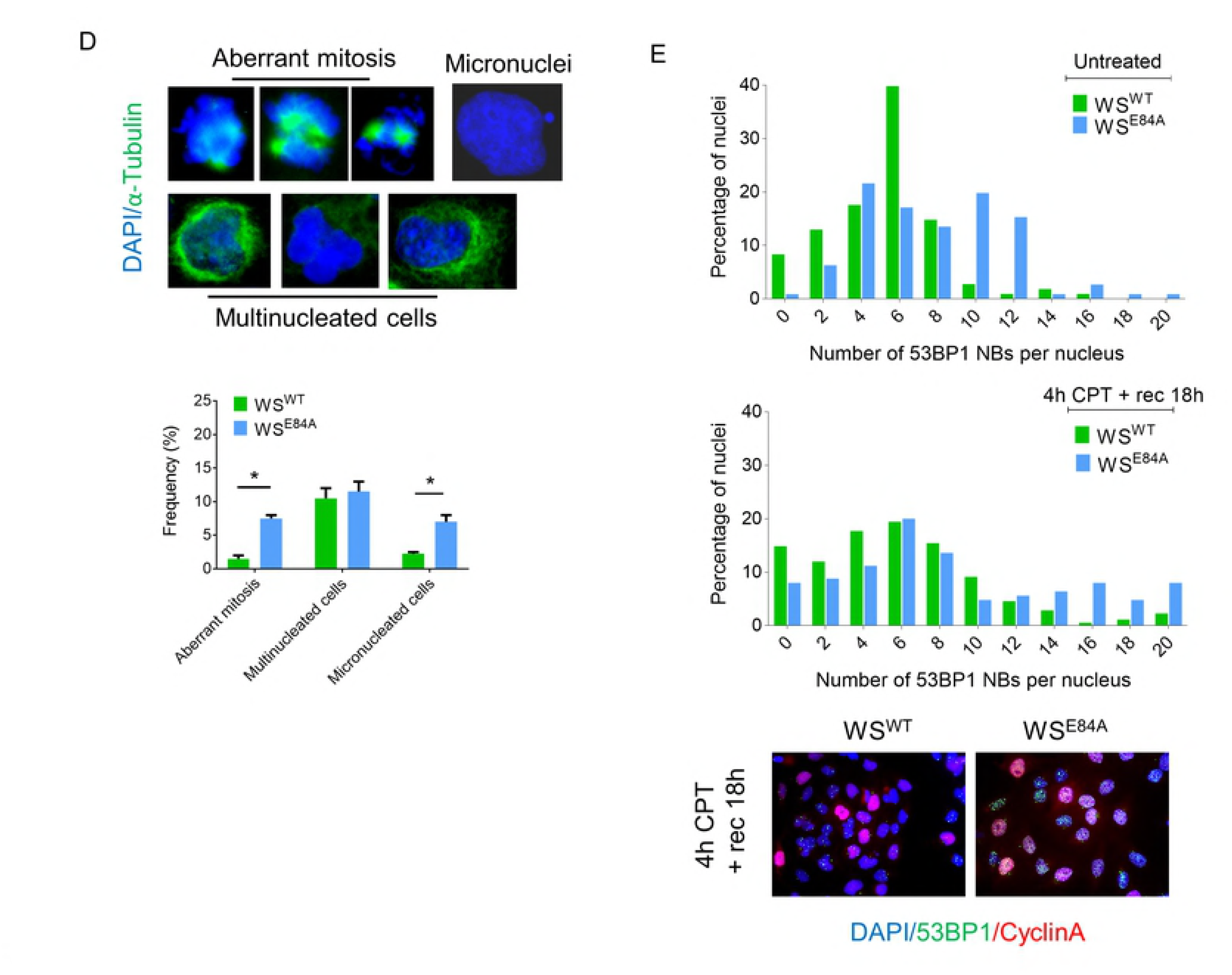
WRN exonuclease-deficient cells show enhanced MUS81 phosphorylation on S87 and mitotic defects. (A) Anti-pS87MUS81 immunofluorescence staining (red) was performed in wild-type and WRN exonuclease-dead expressing cells. The S-phase cells (green) were revealed with short EdU pulse followed by Click-IT reaction. Nuclei were depicted with DAPI staining (blue). The mean frequency (±SE; n=3) of pS87-MUS81-positive nuclei are indicated in the representative images. (B) WS cells expressing the WS^E84A^ mutant were treated with CPT for 4h and then released in fresh medium for 18 hours. The RAD51 inhibitor B02 was added with CPT and during the recovery. The frequency (± SE; n=2) of pS87-MUS81-positive nuclei are indicated as percentage in the representative images. (C) The graph shows the mean percentage ± SE of bulky anaphase bridges analysed in untreated and CPT-treated cells expressing WRN wild-type and the exonuclease-deficient mutant. The number of anaphases counted for each experimental point are indicated above as *n*. Randomly-selected representative anaphases with bridges are shown. (D) Representative images of mitotic aberrations analysed in α-Tubulin (green) and DAPI-stained cells are shown above the graph indicating the frequency of each event after treatment with 50nM of CPT for 4h followed by a 18h recovery. Data are presented as mean±SE. Statistical analysis was performed by the ANOVA test (\**P <* 0.5). (E) Analysis of 53BP1 NBs. Cells were either untreated or treated with 50nM CPT as indicated. Samples were subjected to immunofluorescence using anti-53BP1 and anti-Cyclin A to evaluate 53BP1 fluorescence only in G1 cells (Cyclin A-negative). Graphs show the frequency of each class of nuclei in two independent replicates. Representative images from CPT-treated samples are shown.

Enhanced activation of MUS81 in mitosis in WS^E84A^ cells might be indicative of persistence of unresolved DNA intermediates, which could induce mitotic abnormalities or segregation defects. To assess if loss of WRN exonuclease during the response to low-dose of CPT could result in segregation defects, we first analysed the presence of bulky anaphase bridges in DAPI-stained cells (Fig. 7C). Interestingly, anaphase cells were highly enriched in WRN exonuclease-deficient cells (Fig. 7C; see numbers above the bars), suggesting that these cells may have a delayed exit from anaphase. Of note, the number of anaphases with bridges was very low in WRN exonuclease-deficient cells as compared to cells expressing WRN wild-type (Fig. 7C). Since delayed exit from anaphase might derive from mitotic defects and could result in post-mitotic abnormalities, we decided to evaluate the presence of aberrant mitoses, multinucleated cells and micronuclei. As shown in Figure 7D, we found that both aberrant mitosis and cells with micronuclei were increased by loss of WRN exonuclease activity. In contrast, no difference in multinucleated cells was found between cells expressing wild-type or exo-dead WRN (Fig. 7D).

The presence of under-replicated DNA or the persistence of unresolved DNA intermediates in G2/M triggers the formation of 53BP1 NBs in the subsequent G1 phase [33]. Since WRN exonuclease-deficient cells showed persistence of under-replicated DNA, we investigated whether they accumulated 53BP1 NBs. To this end, we performed immunofluorescence against 53BP1 and CyclinA in cells after recovery from 4h treatment with CPT and scored the number of 53BP1 NBs-positive cells in the CyclinA-negative population (i.e. G1 cells). As shown in Figure 7E, the number of 53BP1 NBs in WS^E84A^ cells was higher than in wild-type cells even in untreated conditions. Treatment with CPT enhanced the number of 53BP1 NBs in wild-type and in WRN exonuclease-deficient cells; however, the increase was substantially higher in cells expressing the exo-dead WRN (Fig. 7E and Fig. S5).

Notably, inhibition of RAD51 in WRN exonuclease-deficient cells resulted in persistent accumulation of mitotic cells in pro-metaphase (Fig. S4). This result suggested that loss of RAD51 undermines correct mitotic progression in WRN-exonuclease-deficient cells but prevented assessment of any correlation between enhanced engagement of RAD51 and mitotic defects or formation of 53BP1 NBs.

As in WRN exonuclease-deficient cells engagement of RAD51 is functionally related to elevated levels of S87-MUS81 phosphorylation (Fig. 7B), we next analysed whether inactivation of the mitotic function of MUS81 by overexpression of the unphosphorylable S87A-MUS81 mutant (ref) in these cells could aggravate the mitotic defects. Interestingly, over-expression of the S87A-MUS81 mutant in wild-type cells did not affect the percentage of anaphase bridges and micronuclei, while it increased the number of multinucleated cells (Fig. 8A). In sharp contrast, expression of the S87A-MUS81 mutant substantially aggravated the mitotic defects in WRN exonuclease-deficient cells (Fig. 7A). Indeed, expression of the S87A-MUS81 protein increased the number of anaphase bridges and micronucleated cells. Similarly, expression of the S87A-MUS81 mutant enhanced the presence of 53BP1 NBs in WS^E84A^ cells (Fig. 8B).

**Figure 8.**
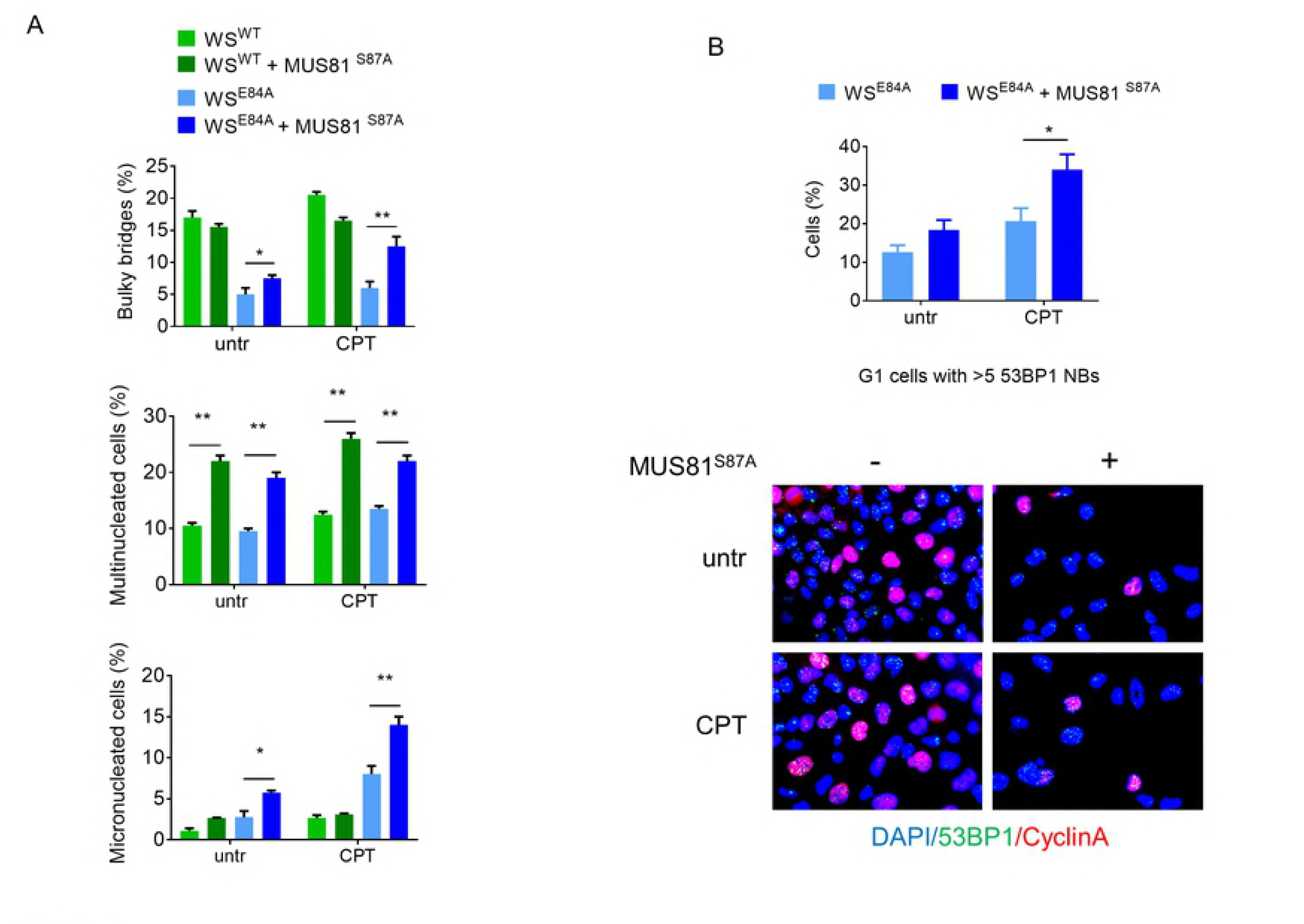
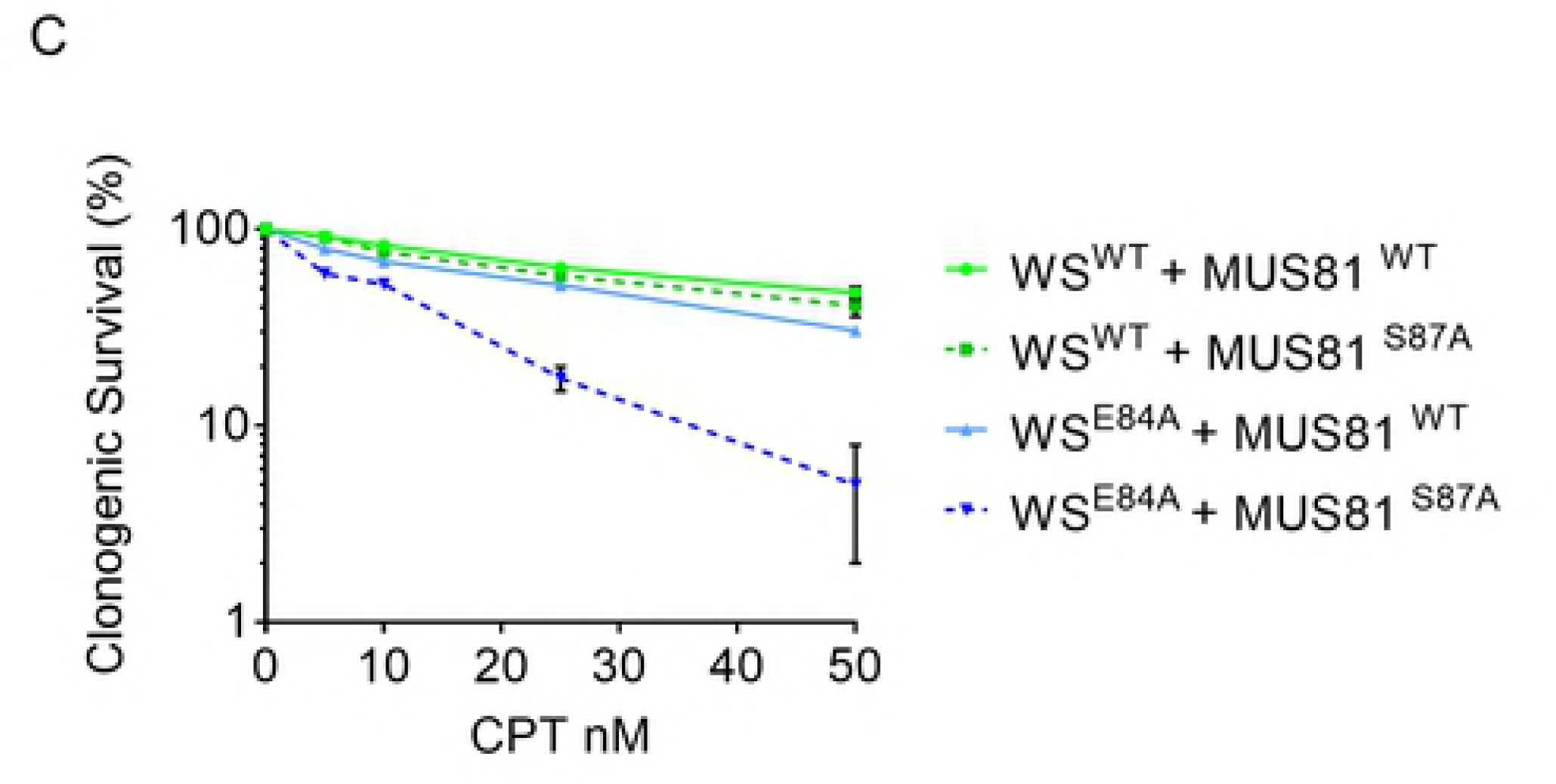
MUS81^S87A^ mutant overexpression aggravates the mitotic phenotypes of WRN exonuclease-deficient cells. (A) Bulky bridges, multinucleated and micro-nucleated cells were analysed with or without MUS81^S87A^ mutant overexpression, in WRN wild-type and exonuclease-deficient cells. The graph represents the frequency of the aberration analysed in two independent experiments ± SE. (B) Analysis of 53BP1 NBs formation in Cyclin A-negative cells. Representative images of fluorescence cells stained with anti-53BP1 (green) and Cyclin A (red). Nuclear DNA was counterstained with DAPI (blue). For each point at least 300 nuclei were counted and cells with > 5 53BP1 NBs were considered as positive. The graph shows the quantification of 53BP1-positive G1 cells. (C) Clonogenic assay in cells treated with low-doses of CPT. Cells were exposed to different doses of CPT for 18h, re-plated at low density and survival evaluated as percentages of colonises normalised against the untreated. Statistical analyses in A-C were performed by ANOVA test (* *P* < 0.5; ** *P* < 0.1).

To further assess the biological significance of the MUS81 hyperactivation observed in mitosis in the absence of the WRN exonuclease, we evaluated the sensitivity to nanomolar doses of CPT by clonogenic survival. As shown in Figure 8C, WRN exonuclease-deficient cells were slightly more sensitive to CPT then wild-type cells. Overexpression of the wild-type MUS81 resulted in a mild increase in sensitivity in wild-type cells but not in cells expressing the exo-dead WRN (Figure 7C). In contrast, overexpression of the S87A-MUS81 resulted in a substantial increase in the sensitivity of WRN exonuclease-deficient cells to CPT (Fig. 8C).

Altogether, our data indicate that the enhanced engagement of RAD51 observed in the absence of the WRN exonuclease requires the increased activation of the MUS81 complex in mitosis. Therefore, expression of a MUS81 mutant that disables mitotic activation of the MUS81/EME1 complex increases mitotic abnormalities and sensitivity to CPT of WRN exonuclease-deficient cells. Thus, in the absence of the WRN exonuclease, hyperactivation of the MUS81 complex functions as a fail-safe system that maintains mitotic abnormalities at low levels, allowing survival.

## DISCUSSION

In recent years, an increasing interest arose around alternative mechanisms of fork processing and fork degradation since they correlate with response to chemotherapeutics in cells that are deficient for the primary pathway(s) as described in the absence of BRCA1/2 [15,17]. Most of these studies focused on the early events occurring in the absence of BRCA1 or BRCA2, but few of them investigated mechanisms involved during recovery from replication stress [29,34]. Furthermore, loss of BRCA1 or BRCA2 affects recombination as well as fork protection and this prevents the investigation of the role of recombination for the recovery of replication forks undergoing degradation. Recently, we reported that the WRN exonuclease activity protects against fork degradation when cells are treated with clinically-relevant doses of CPT [11]. Here, we used WRN exonuclease-deficient cells as a model to assess what happens at destabilised perturbed forks when treatment with nanomolar doses of CPT is prolonged. We find that, in the absence of the WRN exonuclease, nascent strands undergo continuous degradation that produces accumulation of ssDNA. This late accumulation of ssDNA follows its disappearance at early time points because of the activities of MRE11 and EXO1 [11]. Interestingly, the late wave of ssDNA at perturbed forks is only minimally affected by inactivation of each single exonuclease acting at perturbed forks, and is only reduced when MRE11 and DNA2 are both inhibited. This suggests that multiple nucleases take over with time at forks destabilised by the absence of WRN exonuclease while most of the degradation observed in cells deficient of BRCA1/2, or other factors assisting RAD51, seems to involve only MRE11-EXO1 [35–38]. Treatment with nanomolar doses of CPT does not induce DSBs unless treatment is prolonged [12,13]. Interestingly, loss of the WRN exonuclease makes cells resistant to the induction of DSBs after prolonged treatment with nanomolar CPT. Induction of DSBs in response to nanomolar doses of CPT has been correlated with activation of RECQ1 possibly to promote restart of those forks that failed to be processed otherwise [13]. Loss of the ability to induce DSBs at forks would be consistent with engagement of a distinct fork recovery mechanism in cells expressing the exo-dead WRN protein. Indeed, WRN exonuclease-deficient cells do not show RECQ1 PARylation [11], which is required to avoid unscheduled RECQ1 activation [13], which, together with the absence of DSBs support a pathway switch at CPT-perturbed forks.

Consistent with the pathway switch, inhibition of MRE11 is sufficient to restore DSBs after prolonged treatment with a low-dose of CPT in WRN exonuclease-deficient cells, suggesting that formation of DSBs does not necessarily occur downstream of pathological fork processing. Interestingly, RAD51 is strongly accumulated in the absence of the WRN exonuclease and persists during recovery from treatment. An elevated engagement of RAD51 in the absence of the WRN exonuclease has been reported in *Drosophila* [39], suggesting that the role of WRN exonuclease at perturbed forks is conserved. Similarly, unscheduled exonuclease-mediated processing of perturbed forks in yeast has been recently shown to engage a RAD51-mediated pathway [40]. Furthermore, parental ssDNA, a readout of template gaps, also accumulates in the absence of WRN exonuclease. RAD51 binds to ssDNA and initiates recombination [19,25]. The concomitant accumulation of ssDNA and elevated recruitment of RAD51 in the absence of the WRN exonuclease would be consistent with the engagement of gaps left behind inactivated forks in a template-switch mode of replication recovery, as shown after DNA damage in *Xenopus* egg extracts [23]. Consistent with this possibility, WRN exonuclease-deficient cells show enhanced inactivation of CPT-perturbed forks and new origin firing. Moreover, although RAD51 has been implicated in fork restart [24,41], in WRN exonuclease-deficient cells, RAD51 is not involved in fork reactivation after CPT treatment. Indeed, its inhibition does not increase fork inactivation. This result is in agreement with the participation of RAD51 in “gap repair” and supports the notion that prolonged treatment with nanomolar CPT channels perturbed forks into alternative fork processing pathways if the function of WRN exonuclease is absent. Consistent with this, in wild-type cells, inhibition of RAD51 reduces fork reactivation. This is not unexpected since RAD51 plays crucial roles in both fork remodelling and stability [14,16]. In addition, as RAD51 and WRN have been proposed to cooperate during recovery from fork arrest [42], it is reasonable to speculate that loss of WRN function also compromises the normal activity of RAD51 at fork.

Loss of WRN exonuclease results in a mild defect in the activation of CHK1. Activation of the ATR-dependent checkpoint requires formation of ssDNA [43–45]. In the absence of the WRN exonuclease ssDNA accumulates but is hijacked by RAD51 and it is not completely free for the binding of checkpoint factors. Indeed, in WRN exonuclease-deficient cells, TopBP1 and its binding factor RAD9 are not more highly associated with ssDNA as compared with the wild-type. Notably, phosphorylation of ATR, a readout of its activation, is indistinguishable from the wild-type. It suggests that ATR also gets activated independently from ssDNA. Alternatively, a hyperactivation of ATR is also prevented by sequestering of ssDNA by RAD51. Indeed, overexpression of RAD51 has been shown to affect checkpoint activation [46]. Notably, bypassing of the CHK1 activation defect by expression of a phosphomimetic CHK1 mutant in WRN exonuclease-deficient cells restores normal levels of ssDNA. This suggests that accumulation of ssDNA is also unleashed by reduced CHK1 activation through a positive feedback loop.

The observed elevated recruitment of RAD51, which is used during recovery in the absence of the WRN exonuclease to deal with under-replicated DNA, also leads to elevated phosphorylation of MUS81 at S87. Phosphorylation of MUS81 at S87 occurs in G2/M and is related to resolution of recombination intermediates [32]. Consistently, inhibition of RAD51 reduces S87 phosphorylation in WRN exonuclease-deficient cells. Thus, engagement of RAD51-dependent fork recovery, possibly by template switch since DSBs do not form, results in an increased number of interlinked intermediates calling for resolution by the MUS81 complex. Our data indicate that activation of MUS81 complex in G2/M is essential to overcome segregation defects arising from excessive RAD51-dependent recombination and support proliferation upon treatment with CPT. Indeed, expression of the unphosphorylable S87A-MUS81 mutant increases abnormal mitosis and sensitizes WRN exonuclease-deficient.

Loss of the WRN exonuclease although resulting in fork degradation does not induce MUS81 activation in S-phase, which is observed in the absence of BRCA2 [37]. However, the persistence of under-replicated DNA and requirement of MUS81 complex activity in G2/M shown by WRN exonuclease-deficient cells are also characteristic of BRCA2-deficient cells [34]. Thus, it is tempting to speculate that elevated fork degradation correlates with inability to replicate all the genome. Notably, BRCA2-deficient cells show much more severe mitotic defects [29,34]. In WRN exonuclease-deficient cells, mitotic abnormalities are increased by disabling MUS81 function in mitosis but are likely increased also by impairing RAD51 function during recovery, since RAD51 inhibition results in increased accumulation of parental ssDNA and induces a significant mitotic block. As BRCA2 deficiency also interferes with the post-replicative function of RAD51 [23] it is tempting to speculate that the elevated mitotic defects might be the end-result of combined fork deprotection and recombination defects. Indeed, it is recently demonstrated that mitotic abnormalities in BRCA2-deficient cells are primarily linked to loss of the recombination function of RAD51 using separation-of-function mutants [29].

Collectively, our data show that WRN exonuclease-deficient cells can be a useful model to investigate the fate of deprotected or destabilised replication forks under a clinically-relevant, specific type of replication stress; and, together with published data, they can be summarised in the model shown in Figure 9. In response to nanomolar CPT, perturbed replication forks rapidly undergo fork reversal [12]. The WRN exonuclease is required somehow at this stage to prevent MRE11-dependent degradation. In wild-type cells, with time, reversed forks degenerate into DSBs, possibly because of unscheduled RECQ1-mediated fork restoration [13]. In the absence of WRN exonuclease, perturbed replication forks undergo a further cycle of degradation of nascent strand by MRE11 and/or DNA2, which leads to ssDNA accumulation and engagement of RAD51. The accumulation of ssDNA and possibly engagement of RAD51 make perturbed replication forks resistant to DSBs and interfere with checkpoint signalling, resulting in a mild defect in CHK1 activation. In the absence of WRN exonuclease, more perturbed forks become inactivated over-time and RAD51 is required also during recovery from CPT to support repair at template gaps left behind the inactive forks. Engagement of RAD51 during recovery results in elevated activation of the MUS81 complex in G2/M to deal with intermediates, and to limit mitotic defects and cell death. As CPT is a chemotherapeutic, our data also indicate that tumors with impaired function of the WRN exonuclease can be sensitized to treatment by genetic or chemical interference with the MUS81 complex in mitosis, which is less relevant for survival in cells expressing the WRN wild-type.

**Figure 9.**
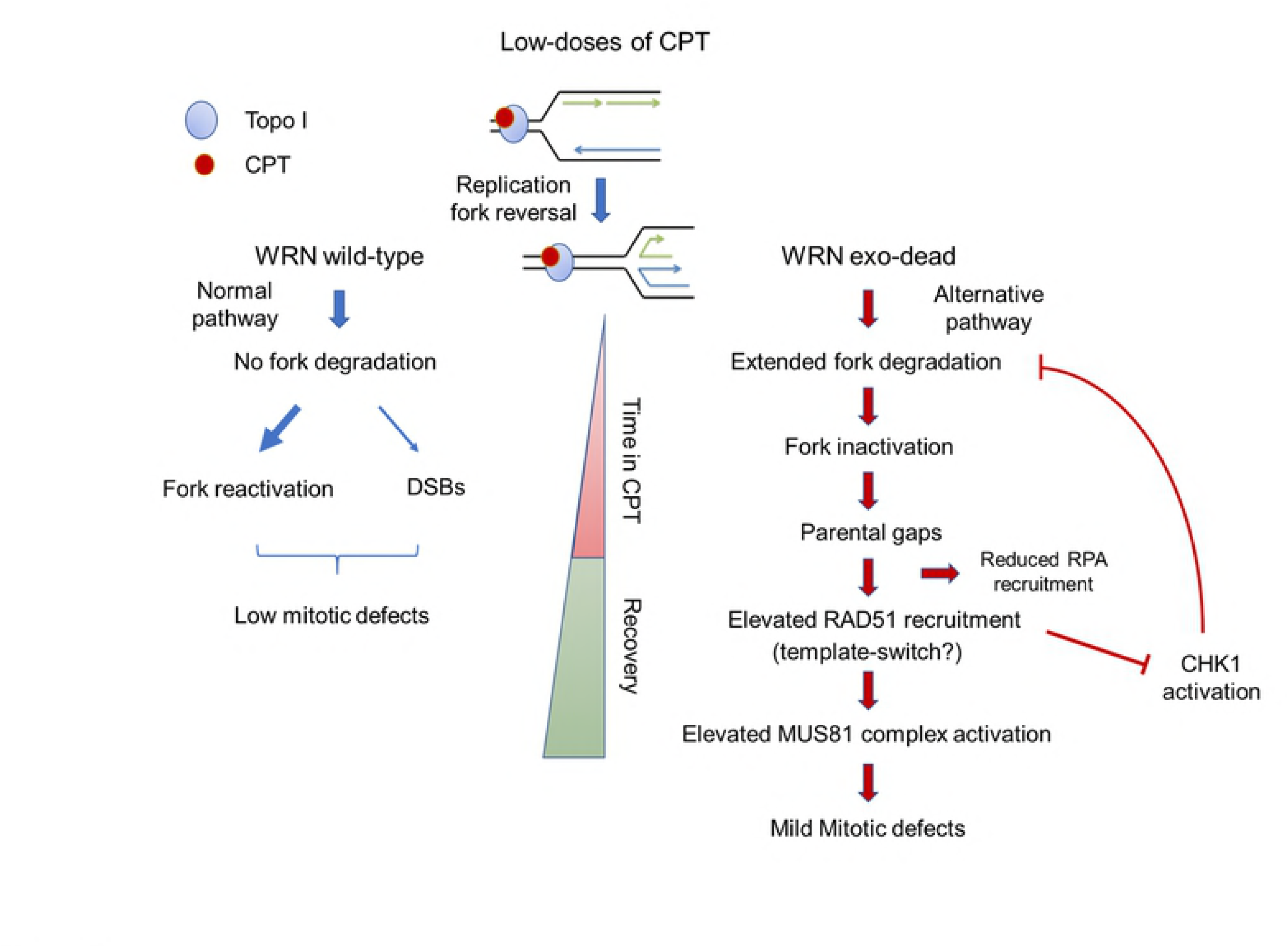
Proposed model of the effect of prolonged treatment with nanomolar CPT doses in absence of WRN exonuclease. (see text for details).

## MATERIALS AND METHODS

### Cell lines and culture conditions

The SV40-transformed WRN-deficient fibroblast cell line (AG11395) was obtained from Coriell Cell Repositories (Camden, NJ, USA). To produce stable cell lines, AG11395 (WS) fibroblasts were transduced with retroviruses expressing the full-length cDNA encoding wild-type WRN (WS^WT^), exonuclease-dead (WS^E84A^), or helicase-dead (WS^K577M^)[47]. All the cell lines were maintained in Dulbecco’s modified Eagle’s medium (DMEM; Life Technologies) supplemented with 10% FBS (Boehringer Mannheim) and incubated at 37 °C in a humidified 5% CO_2_ atmosphere.

### Chemicals

Camptothecin (ENZO Lifesciences) was dissolved in DMSO and a stock solution (10 mM) was prepared and stored at −20°C. Mirin (Calbiochem), an inhibitor of MRE11 exonuclease activity, was used at 50 μM; the B02 compound (Selleck), an inhibitor of RAD51 activity, was used at 27 μM. C5 (ref), DNA2 inhibitor C5 was dissolved in DMSO and used at final concentration of 300 μM [20]. IdU and CldU (Sigma-Aldrich) were dissolved in sterile DMEM at 2.5mM and 200mM respectively and stored at −20°C.

### Plasmids transfection

Plasmid expressing the phospho - mimic (Flag-CHK1^317/345D^) mutant form of CHK1, a kind gift from Professor K.K. Khanna (Queensland Institute of Medical Research, Australia) was generated as described [28]. To express the plasmids, cells were transfected using the Neon^TM^ Transfection System Kit (Invitrogen), according to the manufacturer’s instructions.

### Immunofluorescence assays

Cells were grown on 35-mm coverslips and harvested at the indicated times after treatments. For RAD51 IF, after further washing with PBS, cells were pre-extracted with 0,5% TritonX-100 and fixed with 3% PFA / 2% sucrose at RT for 10min. After blocking in 3% BSA for 15 min, staining was performed with rabbit monoclonal anti-RAD51 (Bioss, 1:100) diluted in a 1% BSA / 0,1% saponin in PBS solution, for 1h at 37° in a humidified chamber. For 53BP1, pS87MUS81 and α-tubulin staining, cells were fixed with 4% PFA at RT for 10 min. Cells were subsequently permeabilized with 0,4% Triton-X100. Staining with primary antibodies diluted in a 1% BSA / 0,1% saponin in PBS solution was carried out for 1h at RT. After extensive washing with PBS, specie-specific fluorophore-conjugated antibody (Invitrogen) was applied for 1h at RT followed by counterstaining with 0.5 mg/ml DAPI. Secondary antibody was used at 1:200 dilution. Images were acquired as greyscale files using Metaview software (MDS Analytical Technologies) and processed using Adobe Photoshop CS3 (Adobe). For each time point, at least 200 nuclei were examined, and foci were scored at 40×. Only nuclei with > 5 foci were considered as positive and were quantified using ImageJ.

### EdU incorporation assay

To label replicated DNA, cells were incubated with 10 μM EdU for 30 minutes. Samples were fixed with 4% PFA at RT for 10 min and cells were subsequently permeabilized with 0,5% Triton-X100. EdU incorporation was detected using the Click-It Edu Alexa Fluor 488 Imaging Kit (Invitrogen) according to the manufacturer’s instructions.

### Antibodies

The primary antibodies used were: anti-pS10H3 (1:1000, Santa Cruz Biotechnologies), anti-Cyclin A (IF: 1:100, Santa Cruz Biotechnologies), anti-53BP1 (1:300, Millipore), anti-BrdU (1:80, Abcam; CldU detection), anti-BrdU (1:50, Becton Dickinson; anti-IdU detection), anti-pS87MUS81 (ref 1:200), anti-RAD51 (1:1000, Bioss Antibodies), anti-αTubulin (1:50, Sigma-Aldrich) and anti-Lamin B1 (1:10000, Abcam).

### Chromatin fractionation and Western blot analysis

Chromatin fractionation experiments were performed as previously described [48]. Western blotting was performed using standard methods. Blots were incubated with primary antibodies against: rabbit anti-pCHK1(S345) (Cell Signalling Technology), mouse anti-CHK1 (Santa Cruz Biotechnology), rabbit anti-RAD51 (Bioss Antibodies), mouse anti-RPA32 (Calbiochem), rabbit anti-RPA70 (GeneTex), mouse anti-GAPDH (Santa Cruz Biotechnology) and rabbit anti-Lamin B1 (Abcam). After incubations with horseradish peroxidase-linked secondary antibodies (1:20000, Jackson Immunosciences), the blots were developed using the chemiluminescence detection kit WesternBright ECL HRP substrate (Advansta) according to the manufacturer’s instructions. Quantification was performed on scanned images of blots using the Image Lab software, and the values shown on the graphs represent normalization of the protein content evaluated through LaminB1 or GAPDH immunoblotting.

### Clonogenic survival

Cells were plated onto 35mm dishes, after 24h they were treated with different doses of CPT. After 18, cells were washed, trypsinized and seeded in 60mm dishes. After 14-21 days, plates were stained with crystal violet and colonies counted.

### DNA fibres analysis

DNA fibres were prepared, spread out and immunodecorated as previously described [11]. Images were acquired randomly from fields with untangled fibres using Eclipse 80i Nikon Fluorescence Microscope, equipped with a VideoConfocal (ViCo) system. The length of labeled tracks were measured using the Image-Pro-Plus 6.0 software. A minimum of 100 individual fibres were analysed for each experiment and the mean of at least three independent experiments presented.

### Detection of nascent single-stranded DNA

To detect nascent single-stranded DNA (ssDNA), cells were plated onto 22×22 coverslips in 35mm dishes. After 24h, the cells were labelled for 15 min before the treatment with 250μM IdU (Sigma-Aldrich), cells were then treated with CPT 5μM for different time points. Next, cells were washed with PBS, permeabilized with 0.5% Triton X-100 for 10 min at 4°C and fixed wit 2% sucrose, 3% PFA. For ssDNA detection, cells were incubated with primary mouse anti-BrdU antibody (Becton Dickinson) for 1h at 37°C in 1%BSA/PBS, followed by Alexa Fluor488-conjugated goat-anti-Mouse (Invitrogen), and counterstained with 0.5μg/ml DAPI. Slides were analysed with Eclipse 80i Nikon Fluorescence Microscope, equipped with a VideoConfocal (ViCo) system. For each time point, at least 100 nuclei were scored at 60×. Parallel samples either incubated with the appropriate normal serum or only with the secondary antibody confirmed that the observed fluorescence pattern was not attributable to artefacts. Fluorescence intensity for each sample was then analysed using ImageJ software.

### Statistical analysis

All the data are presented as means of at least two independent experiments. Statistical comparisons of WS^WT^ or WRN-mutant cells to their relevant control were analysed by ANOVA or Mann-Whitney test. P < 0.5 was considered as significant.

## ACKNOWLEDGMENTS

We are grateful to Prof. Massimo Lopes (IMCR, University of Zurich) for scientific discussion. We thank all members of our laboratories for discussion. This work was supported by Associazione Italiana per la Ricerca sul Cancro (AIRC) to PP (IG17383) and to AF (IG119971), and in part by NIH R01CA085344 to BHS, R50CA211397 to LZ and GM123554 to JLC.

## AUTHOR CONTRIBUTIONS

F.A.A. performed the analysis of CHK1 phosphorylation, fork recruitment by PLA and chromatin fractionation, and performed experiments to determine DNA damage. A.P. performed the analysis of MUS81 phosphorylation and experiments to evaluate mitotic abnormalities. E.M. analysed the persistence of RAD51 and parental ssDNA. F.A.A., A.P., E.M. analysed data, contributed to designing the experiments and writing the manuscript. A.F. and P.P. designed experiments, analysed data and wrote the paper. L.Z., J.L.C. and B.H.S. provided the DNA2 inhibitor C5, advised the relevant experiments, and revised the manuscript. All authors approved the paper.

## SUPPORTING INFORMATION LEGENDS

**Supplementary Figures and Legends.** The file contains five supplementary figures and their legends.

## REFERENCES

1. Técher H, Koundrioukoff S, Nicolas A, Debatisse M. The impact of replication stress on replication dynamics and DNA damage in vertebrate cells. Nat Rev Genet. Nature Publishing Group; 2017; doi:10.1038/nrg.2017.46

2. Franchitto A, Pichierri P. Replication fork recovery and regulation of common fragile sites stability. Cell Mol Life Sci. 2014;71: 4507–4517. doi:10.1007/s00018-014-1718-9

3. Zeman MK, Cimprich KA. Causes and consequences of replication stress. Nat Cell Biol. 2014;16: 2–9. doi:10.1038/ncb2897

4. Macheret M, Halazonetis TD. DNA Replication Stress as a Hallmark of Cancer. Annu Rev Pathol Mech Dis. 2015;10. doi:10.1146/annurev-pathol-012414-040424

5. Magdalou I, Lopez BS, Pasero P, Lambert S a E. The causes of replication stress and their consequences on genome stability and cell fate. Semin Cell Dev Biol. Elsevier Ltd; 2014;30: 154–164. doi:10.1016/j.semcdb.2014.04.035

6. Hills SA, Diffley JFX. DNA Replication and Oncogene-Induced Replicative Stress. Curr Biol. Elsevier Ltd; 2014;24: R435–R444. doi:10.1016/j.cub.2014.04.012

7. Ciccia A, Elledge SJ. The DNA Damage Response: Making It Safe to Play with Knives. Mol Cell. Elsevier Inc.; 2010;40: 179–204. doi:10.1016/j.molcel.2010.09.019

8. Pichierri P, Ammazzalorso F, Bignami M, Franchitto A. The Werner Syndrome protein: Linking the replication checkpoint response to genome stability. Aging (Albany NY). 2011;3: 311–318. doi:100293 [pii]

9. Rossi ML, Ghosh AK, Bohr V a. Roles of Werner syndrome protein in protection of genome integrity. DNA Repair (Amst). Elsevier B.V.; 2010;9: 331–44. doi:10.1016/j.dnarep.2009.12.011

10. Franchitto A, Pichierri P. Understanding the molecular basis of common fragile sites instability: Role of the proteins involved in the recovery of stalled replication forks. Cell Cycle. 2011;10: 4039–4046. doi:10.4161/cc.10.23.18409

11. Iannascoli C, Palermo V, Murfuni I, Franchitto A, Pichierri P. The WRN exonuclease domain protects nascent strands from pathological MRE11/EXO1-dependent degradation. Nucleic Acids Res. 2015;43: 9788–803. doi:10.1093/nar/gkv836

12. Ray Chaudhuri A, Hashimoto Y, Herrador R, Neelsen KJ, Fachinetti D, Bermejo R, et al. Topoisomerase I poisoning results in PARP-mediated replication fork reversal. Nature Structural & Molecular Biology. Nature Publishing Group; 2012. pp. 417–423. doi:10.1038/nsmb.2258

13. Berti M, Ray Chaudhuri A, Thangavel S, Gomathinayagam S, Kenig S, Vujanovic M, et al. Human RECQ1 promotes restart of replication forks reversed by DNA topoisomerase I inhibition. Nat Struct Mol Biol. Nature Publishing Group; 2013;20: 347–54. doi:10.1038/nsmb.2501

14. Bhat KP, Cortez D. RPA and RAD51: fork reversal, fork protection, and genome stability. Nat Struct Mol Biol. 2018;25: 446–453. doi:10.1038/s41594-018-0075-z

15. Quinet A, Lemaçon D, Vindigni A. Replication Fork Reversal: Players and Guardians. Mol Cell. 2017;68: 830–833. doi:10.1016/j.molcel.2017.11.022

16. Kolinjivadi AM, Sannino V, de Antoni A, Técher H, Baldi G, Costanzo V. Moonlighting at replication forks - a new life for homologous recombination proteins BRCA1, BRCA2 and RAD51. FEBS Lett. 2017; 1–18. doi:10.1002/1873-3468.12556

17. Feng W, Jasin M. Homologous Recombination and Replication Fork Protection: BRCA2 and More! Cold Spring Harb Symp Quant Biol. 2017;82: 329–338. doi:10.1101/sqb.2017.82.035006

18. Costanzo V. Brca2, Rad51 and Mre11: Performing balancing acts on replication forks. DNA Repair (Amst). Elsevier B.V.; 2011;10: 1060–1065. doi:10.1016/j.dnarep.2011.07.009

19. Pellegrini L, Venkitaraman A. Emerging functions of BRCA2 in DNA recombination. Trends Biochem Sci. 2004;29: 310–316. doi:10.1016/j.tibs.2004.04.009

20. Liu W, Zhou M, Li Z, Li H, Polaczek P, Dai H, et al. A Selective Small Molecule DNA2 Inhibitor for Sensitization of Human Cancer Cells to Chemotherapy. EBioMedicine. The Authors; 2016;6: 73–86. doi:10.1016/j.ebiom.2016.02.043

21. Murfuni I, Nicolai S, Baldari S, Crescenzi M, Bignami M, Franchitto a, et al. The WRN and MUS81 proteins limit cell death and genome instability following oncogene activation. Oncogene. Nature Publishing Group; 2012;32: 610–20. doi:10.1038/onc.2012.80

22. Hanada K, Budzowska M, Davies SL, van Drunen E, Onizawa H, Beverloo HB, et al. The structure-specific endonuclease Mus81 contributes to replication restart by generating double-strand DNA breaks. Nat Struct Mol Biol. 2007;14: 1096–1104. doi:10.1038/nsmb1313

23. Hashimoto Y, Chaudhuri AR, Lopes M, Costanzo V. Rad51 protects nascent DNA from Mre11-dependent degradation and promotes continuous DNA synthesis. Nat Struct Mol Biol. 2010;17: 1305–1311. doi:10.1038/nsmb.1927

24. Petermann E, Orta ML, Issaeva N, Schultz N, Helleday T. Hydroxyurea-Stalled Replication Forks Become Progressively Inactivated and Require Two Different RAD51-Mediated Pathways for Restart and Repair. Mol Cell. 2010;37: 492–502. doi:10.1016/j.molcel.2010.01.021

25. Carr AM, Lambert S. Replication stress-induced genome instability: The dark side of replication maintenance by homologous recombination. J Mol Biol. Elsevier Ltd; 2013;425: 4733–4744. doi:10.1016/j.jmb.2013.04.023

26. Lee J, Kumagai A, Dunphy WG. The Rad9-Hus1-Rad1 checkpoint clamp regulates interaction of TopBP1 with ATR. J Biol Chem. 2007;282: 28036–28044. doi:10.1074/jbc.M704635200

27. Delacroix S, Wagner JM, Kobayashi M, Yamamoto KI, Karnitz LM. The Rad9-Hus1-Rad1 (9-1-1) clamp activates checkpoint signaling via TopBP1. Genes Dev. 2007;21: 1472–1477. doi:10.1101/gad.1547007

28. Gatei M, Sloper K, Sörensen C, Syljuäsen R, Falck J, Hobson K, et al. Ataxia-telangiectasia-mutated (ATM) and NBS1-dependent phosphorylation of Chk1 on Ser-317 in response to ionizing radiation. J Biol Chem. 2003;278: 14806–14811. doi:10.1074/jbc.M210862200

29. Feng W, Jasin M. BRCA2 suppresses replication stress-induced mitotic and G1 abnormalities through homologous recombination. Nat Commun. 2017;8: 525. doi:10.1038/s41467-017-00634-0

30. Ying S, Minocherhomji S, Chan KL, Palmai-Pallag T, Chu WK, Wass T, et al. MUS81 promotes common fragile site expression. Nat Cell Biol. Nature Publishing Group; 2013;15: 1001–7. doi:10.1038/ncb2773

31. Naim V, Wilhelm T, Debatisse M, Rosselli F. ERCC1 and MUS81-EME1 promote sister chromatid separation by processing late replication intermediates at common fragile sites during mitosis. Nat Cell Biol. Nature Publishing Group; 2013;15: 1008–15. doi:10.1038/ncb2793

32. Palma A, Pugliese GM, Murfuni I, Marabitti V, Malacaria E, Rinalducci S, et al. Phosphorylation by CK2 regulates MUS81/EME1 in mitosis and after replication stress. Nucleic Acids Res. Oxford University Press; 2018;46: 5109–5124. doi:10.1093/nar/gky280

33. Lukas C, Savic V, Bekker-Jensen S, Doil C, Neumann B, Pedersen RS, et al. 53BP1 nuclear bodies form around DNA lesions generated by mitotic transmission of chromosomes under replication stress. Nat Cell Biol. Nature Publishing Group; 2011;13: 243–253. doi:10.1038/ncb2201

34. Lai X, Broderick R, Bergoglio V, Zimmer J, Badie S, Niedzwiedz W, et al. MUS81 nuclease activity is essential for replication stress tolerance and chromosome segregation in BRCA2-deficient cells. Nat Commun. 2017;8: 15983. doi:10.1038/ncomms15983

35. Schlacher K, Wu H, Jasin M. A Distinct Replication Fork Protection Pathway Connects Fanconi Anemia Tumor Suppressors to RAD51-BRCA1/2. Cancer Cell. Elsevier Inc.; 2012;22: 106–116. doi:10.1016/j.ccr.2012.05.015

36. Schlacher K, Christ N, Siaud N, Egashira A, Wu H, Jasin M. Double-Strand Break Repair-Independent Role for BRCA2 in Blocking Stalled Replication Fork Degradation by MRE11. Cell. Elsevier Inc.; 2011;145: 529–542. doi:10.1016/j.cell.2011.03.041

37. Lemaçon D, Jackson J, Quinet A, Brickner JR, Li S, Yazinski S, et al. MRE11 and EXO1 nucleases degrade reversed forks and elicit MUS81-dependent fork rescue in BRCA2-deficient cells. Nat Commun. 2017;8: 860. doi:10.1038/s41467-017-01180-5

38. Kolinjivadi AM, Sannino V, De Antoni A, Zadorozhny K, Kilkenny M, Técher H, et al. Smarcal1-Mediated Fork Reversal Triggers Mre11-Dependent Degradation of Nascent DNA in the Absence of Brca2 and Stable Rad51 Nucleofilaments. Mol Cell. 2017;67: 867–881.e7. doi:10.1016/j.molcel.2017.07.001

39. Bolterstein E, Rivero R, Marquez M, McVey M. The Drosophila Werner exonuclease participates in an exonuclease-independent response to replication stress. Genetics. 2014;197: 643–52. doi:10.1534/genetics.114.164228

40. García-Rodríguez N, Morawska M, Wong RP, Daigaku Y, Ulrich HD. Spatial separation between replisome- and template-induced replication stress signaling. EMBO J. 2018; e98369. doi:10.15252/embj.201798369

41. Hashimoto Y, Puddu F, Costanzo V. RAD51- and MRE11-dependent reassembly of uncoupled CMG helicase complex at collapsed replication forks. Nat Struct Mol Biol. Nature Publishing Group; 2011;19: 17–24. doi:10.1038/nsmb.2177

42. Sidorova JM, Kehrli K, Mao F, Monnat R. Distinct functions of human RECQ helicases WRN and BLM in replication fork recovery and progression after hydroxyurea-induced stalling. DNA Repair (Amst). 2013;12: 128–139. doi:10.1016/j.dnarep.2012.11.005

43. Shechter D, Costanzo V, Gautier J. Regulation of DNA replication by ATR: signaling in response to DNA intermediates. DNA Repair (Amst). 2004;3: 901–8. doi:10.1016/j.dnarep.2004.03.020

44. Friedel AM, Pike BL, Gasser SM. ATR/Mec1: coordinating fork stability and repair. Curr Opin Cell Biol. 2009;21: 237–244. doi:10.1016/j.ceb.2009.01.017

45. Flynn RL, Zou L. ATR: A master conductor of cellular responses to DNA replication stress. Trends Biochem Sci. 2011;36: 133–140. doi:10.1016/j.tibs.2010.09.005

46. Parplys AC, Seelbach JI, Becker S, Behr M, Wrona A, Jend C, et al. High levels of RAD51 perturb DNA replication elongation and cause unscheduled origin firing due to impaired CHK1 activation. Cell Cycle. 2015;14: 3190–202. doi:10.1080/15384101.2015.1055996

47. Pirzio LM, Pichierri P, Bignami M, Franchitto A. Werner syndrome helicase activity is essential in maintaining fragile site stability. J Cell Biol. 2008;180: 305–314. doi:10.1083/jcb.200705126

48. Murfuni I, Basile G, Subramanyam S, Malacaria E, Bignami M, Spies M, et al. Survival of the Replication Checkpoint Deficient Cells Requires MUS81-RAD52 Function.Maizels N, editor. PLoS Genet. 2013;9: e1003910. doi:10.1371/journal.pgen.1003910

